# SS18::SSX redistributes BAF chromatin remodelers selectively to activate and repress transcription

**DOI:** 10.1101/2024.05.14.594253

**Authors:** Jinxiu Li, Li Li, Kyllie Smith-Fry, Zaki F. Wilmot, Lara Carroll, Linda Morrison, Xinyi Ge, Mary Nelson, Lesley A. Hill, Xiaoyang Zhang, Torsten O. Nielsen, Martin Hirst, T. Michael Underhill, Bradley R. Cairns, Kevin B. Jones

**Author notes:** To whom correspondence should be addressed 2000 Circle of Hope Drive, 3726 HCI RS, Salt Lake City, UT 84112.

## Abstract

Synovial sarcoma (SyS) is driven by the expression of a chromosomal translocation-generated SS18::SSX fusion oncoprotein^1,2^. This oncoprotein fuses the SSX carboxy terminal tail that binds ubiquitylated histone H2A (H2AK119ub)^3–6^ to the SS18 component of both canonical (CBAF) and non-canonical (GBAF, GLTSCR1-containing) BAF-family chromatin remodeling complexes^7–11^, wherein SS18::SSX reprograms the epigenetic landscape. Mice that express SS18::SSX develop tumors that faithfully recapitulate human SyS histopathologically and molecularly^12–14^, allowing dissection of transcriptional reprogramming *in vivo*^15,16^. Here, we show that SS18::SSX redistributes GBAF complexes broadly to promoters and distal enhancers decorated by H2AK119ub, which uncharacteristically lack the transcription-silencing trimethylation of H3K27 that otherwise accompanies H2AK119ub. Fusion-GBAF instead associates with H2AK119ub and transcription-enabling H3K4 trimethylation (promoters), monomethylation (distal enhancers), and H3K27 acetylation (both). CBAF with the fusion redistributes away from typical loci, avoids H2AK119ub-bearing regions, and instead narrowly flanks transcription start sites (TSSs), co-distributed with polybromo BAF (PBAF). Accelerated tumorigenesis in our SyS mouse model followed deletion of *Smarcb1* (PBAF and CBAF), *Pbrm1* (PBAF-specific), and *Arid1a* or *Arid1b* (CBAF-specific). While tumors lacking *Arid1a or Arid1b* retained SyS character, loss of *Smarcb1* or *Pbrm1* resulted in tumors lacking SyS features. These findings suggest that SyS transcriptional reprogramming includes both improper GBAF localization and gene activation at H2AK119ub-bearing regulatory regions and improper gene silencing through loss of CBAF.

## Introduction

SyS is a dangerous malignancy of the soft-tissues, primarily affecting adolescents and young adults^17^. Initially characterized by the expression of epithelial cell markers, Sys arises in mesenchymal tissue compartments^18^. SyS was found consistently to associate with a chromosomal translocation between chromosomes 18 and X, which produces one of three specific fusion genes at the junction point, *SS18::SSX1*, *SS18::SSX2*, or *SS18::SSX4*^1^. In mice, expression of this fusion oncogene in mesenchymal progenitors independently initiates tumorigenesis strongly recapitulating human SyS^12,13^. Critical questions have surrounded how exactly the fusion oncoprotein dysregulates BAF chromatin remodeling complexes to generate oncogenic transcription in the cancer cells.

The study of synovial sarcomagenesis in the mouse has provided a tractable and faithful platform for experimentation, testing elements of the biology observed in human SyS^14,19–23^. Although epigenomics assessments of whole tumors add computational complexity from the infiltrating immune, endothelial, and supporting stromal cells, such non-tumor cell populations comprise a small fraction of the overall cells in SyS in humans and mice, permitting robust interrogation of biology with these methods in whole tumors. We therefore began from a foundation comparing single cell transcriptomes from a discrete cell of origin across SyS development^15^. This approach enabled us to generate a catalog of genes associated with sarcomagenesis, assisting in assigning roles to mapped SS18::SSX positions across the genome. Ultimately, having established that SS18::SSX incorporates into both GBAF and CBAF, altering the prevalence of BAF subtypes in SyS cells^14^, we then leveraged these insights to help determine the basic role of BAF complexes in SyS transcriptional reprogramming.

## Results

### SS18::SSX distributes to proximal and distal regulatory elements bearing ubiquitylated histone 2A

In order to evaluate SS18::SSX distribution across chromatin, genome-wide, we performed immunoprecipitation followed by DNA sequencing of chromatin samples (ChIP-seq) from mouse SyS tumors expressing the human *SS18::*SSX2 (herein denoted as *hSS2*) using an antibody specific for the fusion oncoprotein^24^. SyS tumors were generated by expressing the fusion in mouse hindlimb tissues, initiated by TATCre protein injection at 8 days of life (**Extended Data Fig. 1a-b**). Tumors included a range of histomorphologies across the spectrum of SyS (monophasic, biphasic, and poorly differentiated). ChIP-seq was also employed to assess SyS genome profiles of specific chromatin histone marks, and chromatin conformation by HiChIP was evaluated using an antibody against acetylated histone H3 (H3K27ac) to identify enhancer loops for the purpose of assigning distal regulatory elements to specific genes (**Extended Data Fig. 1c-d**).

In our associated paper, we defined three categories of genes determined by comparison of single cell RNA sequencing (scRNA-seq) of SyS mesenchymal progenitors (cells of origin) and fully transformed SyS cells^15^. These categories comprise: 1) genes that transition from no significant expression in progenitors to expression in SyS (sarcomagenesis activated transcription, SAT genes), 2) genes that are expressed in both the mesenchymal progenitors and the transformed SyS cells (maintained active transcription, MAT genes), and 3) genes that transition from active expression in progenitors to no expression in transformed SyS cells (sarcomagenesis silenced transcription, SST genes).

In order to identify direct targets of SS18::SSX among the genes that increase expression with sarcomagenesis, we evaluated each of the promoters in the SAT genes for the fusion oncoprotein ChIP-seq enrichment surrounding the transcription start site (TSS ± 2kB) and the distal anchors of HiChIP loops that have loop anchors within the same proximal locus (**Fig. 1a**). MAT genes served as housekeeping gene controls in these populations of cells. We then profiled all the called SS18::SSX ChIP-seq peaks (q-value < 0.001) that intersected any TSS in the genome for peak width and enrichment score. Two very distinct patterns emerged among promoter associated peaks: broad peaks, among which the SAT genes were included, and narrow peaks, some of which had very strong fusion enrichment, and among which, MAT genes were included (**Fig. 1b**). This immediately suggested that SS18::SSX targets promoters bearing two patterns, with broad promoter binding designating active transcription in genes across the time course of fusion-driven transformation.

**Fig. 1.**
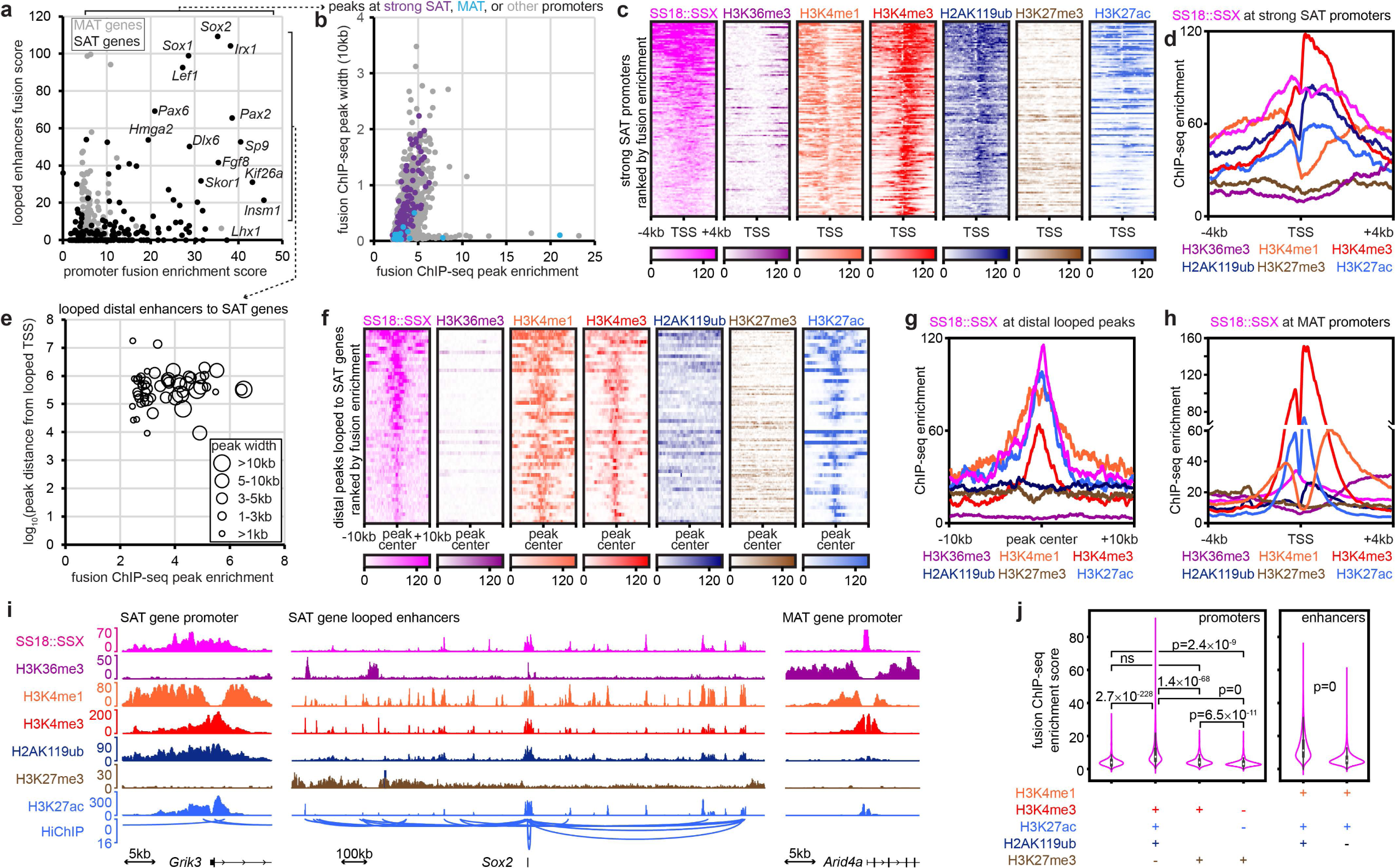
SS18::SSX distributes to the transcriptional regulatory elements of monoubiquitylated histone H2A bearing nucleosomes. (a) Fusion ChIP-seq summed enrichment at enhancer loci (distal anchors ±2kb of H3K27ac-HiChIP loop to selected promoters, with enrichment multiplied by loop score) versus promoter enrichment (transcription start site, TSS±2kb) for genes discretely associated with sarcomagenesis activated transcription (SAT) and maintained active transcription (MAT). (b) Fusion ChIP-seq peak width versus enrichment for all peaks that are within 2kb of any TSS, with those near MAT or strong SAT gene promoters indicated. (c) Heatmaps for ChIP-seq enrichment centered at the TSS of each strong SAT gene promoter. (d) ChIP-seq enrichment plots for histone marks at strong SAT gene promoters. (e) Distal fusion ChIP-seq peak width and distance from its looped SAT gene TSS versus enrichment. (f) Heatmaps for ChIP-seq enrichment centered at the peak center of fusion peaks looped to SAT gene promoters. (g) ChIP-seq enrichment plots at distal fusion peaks looped to SAT gene promoters. (h) ChIP-seq enrichment plots for MAT gene promoters. (i) Example tracks of a histone promoter signature for an SAT gene promoter and looped enhancers, as well as an MAT gene promoter. (j) Violin plots of fusion enrichment at promoters and enhancers with different patterns of intersected peaks called for histone marks or randomly selected promoters across the genome. Any column missing a plus or minus was simply not specified for that ChIP-seq in that category.

In order to characterize the epigenetic attributes of SS18::SSX-occupied SAT gene promoters, we next profiled the distribution of specific histone marks by ChIP-seq (**Fig. 1c-d**, **Extended Data Fig. 1e-f**). These revealed a pattern of strong H2AK119ub and H3K4me3, without H3K27me3, supporting the observation that these typically silent/poised bivalent (H3K4me3- and H3K27me3-marked) developmental gene promoters are converted towards activated transcription, identified in human and mouse SySs^15,16^.

We also sought to characterize the distal (enhancer) binding sites of the fusion that contributed to the regulation of sarcomagenic transcription. Although most loop anchors distant from the promoters of SAT genes colocalized proximally near the TSSs of other genes, others arose from loci distal to all promoters. These distal loci were both broad in width and had strong fusion enrichment (**Fig. 1e**). H2AK119ub was present at a few of these distal loci (**Fig. 1f-g**). However, in striking contrast, the MAT and SST gene promoters had very different patterns of associated histone mark ChIP-seq enrichment, as they lacked significant and strong H2AK119ub enrichment (**Fig. 1h-i**, **Extended Data Fig. 1g-i**).

We next interrogated these signature histone ChIP-seq enrichment patterns across the entire genome to identify other promoters and distal loci that might interact with the fusion with respect to sarcomagenesis. Promoters were therefore defined by the intersection of called peaks (q-value < 0.001) for this SAT pattern of H2K119ub+, H3K4me3+, H3K27ac+, H3K27me3-marks, by the classic bivalent promoter signature (H3K4me3+, H3K27me3+), H3K27me3+ alone promoters, or random promoters across the genome as a control. We also defined distal active enhancers by called intersection peaks for H3K4me1 and H3K27ac, then divided these between those with or without additional called peaks for H2AK119ub. SS18::SSX ChIP-seq enrichment at promoters and distal enhancers that included called peaks for H2AK119ub were significantly stronger than the comparison controls (**Fig. 1j**, **Extended Data Fig. 1j-m**). Thus, histone mark and fusion ChIP-seq enrichment profiles at these SAT pattern-defined promoters and distal enhancers identified strong coincidence of H2AK119ub and the fusion (**Extended Data Fig. 1j-p**), consistent with SSX tail-H2Aub interaction as the dominant mode of GBAF targeting.

### GBAF distributes with the fusion to broad proximal and distal regions; CBAF and PBAF distribute narrowly to active TSSs

The distribution of each BAF family subtype across chromatin was interrogated using ChIP-seq with antibodies against BAF subunits. Single crosslinking was sufficient for BRD9 (a GBAF subunit), SMARCA4 (also known as BRG1, the principal ATPase subunit common to most BAF-family subtypes), and SMARCC1 (also known as BAF155, a subunit incorporating into all BAF-family subtypes), while PBRM1 (a PBAF subunit) and DPF2 (a CBAF subunit) required double cross-linking (**Extended Data Fig. 2a-b**). Strong correlation among biological replicates was confirmed for each antibody. Importantly, strong correlations were found between PBAF and CBAF distributions, as well as between fusion and GBAF distributions (**Fig. 2a**, **Extended Data Fig. 2c**). The stronger correlation of fusion with GBAF than with CBAF (r = 0.73, r = 0.32, Pearson correlation coefficients, respectively) was reflected in strikingly distinct annotation distributions between CBAF/PBAF and GBAF/SS18:SSX. CBAF distributes heavily to promoter regions in SyS, contrasting its distribution in murine stem cells, wherein CBAF predominantly distributes to distal intragenic or intergenic loci (**Fig. 2b**, **Extended Data Fig. 2d**)^25^.

**Fig. 2.**
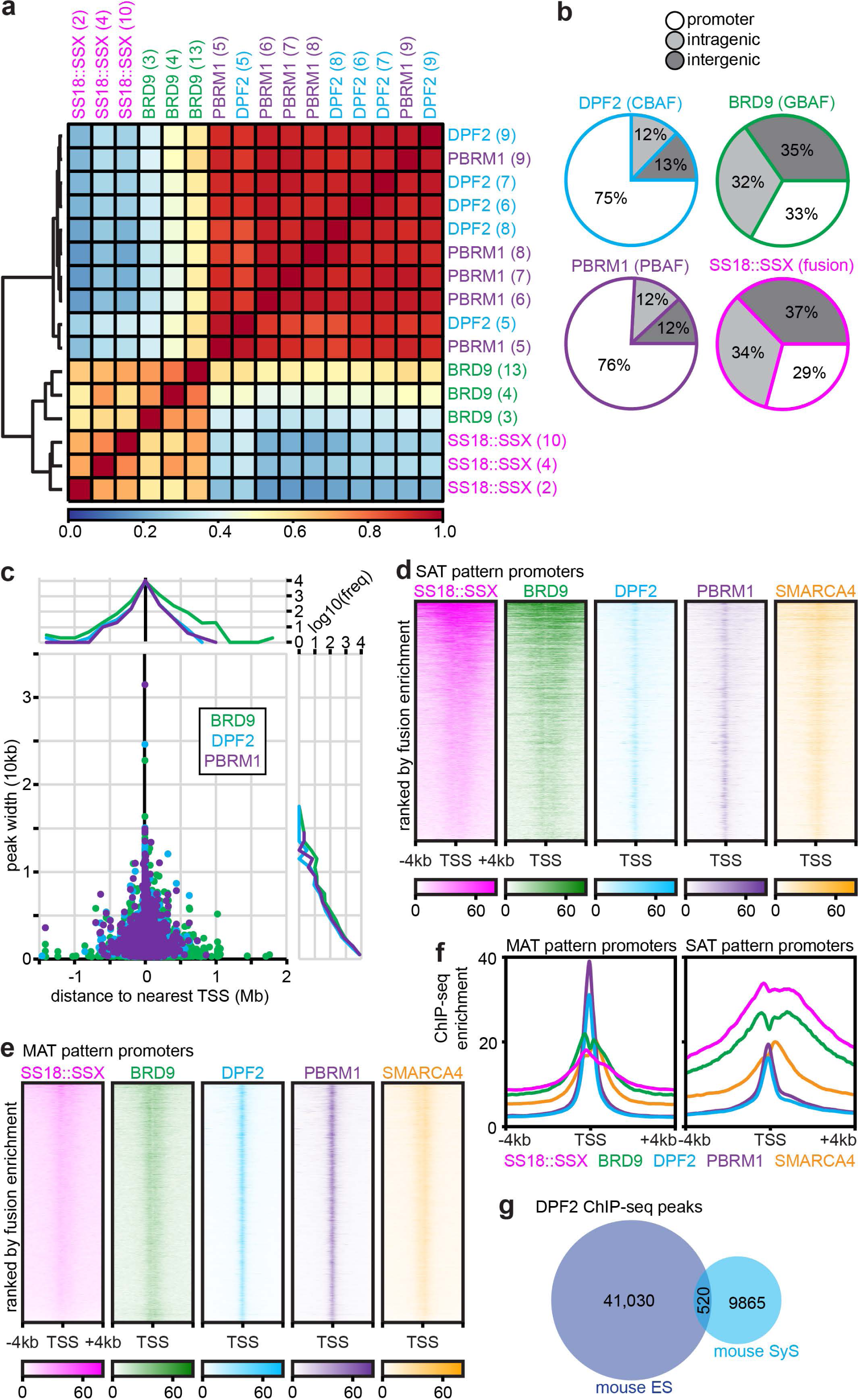
BAF family complex subtypes distribute across chromatin relative to SS18::SSX. (a) Correlation heatmap for genome-wide distributions by ChIP-seq of fusion, BRD9, PBRM1, and DPF2 for individual tumors tested. (b) Chromatin annotation discrepancy between CBAF, PBAF, GBAF, and the fusion. (c) Called peaks width versus distance and histograms from TSS for BRD9, DPF2, and PBRM1. (d) Enrichment heatmaps for SS18::SSX, BRD9. DPF2, PBRM1, and SMARCA4 ChIP-seqs at SAT pattern promoters. (e) Enrichment heatmaps for SS18::SSX, BRD9. DPF2, PBRM1, and SMARCA4 ChIP-seqs at MAT pattern promoters. (f) Enrichment plots for SAT and MAT pattern promoters. (g) Venn diagram of overlap for DPF2 ChIP-seq peaks in mouse embryonic stem cells (ES) per Zhang, et al.^25^, versus mouse synovial sarcomas (SS).

As previously reported, we confirmed a profound loss of CBAF levels in SyS, using glycerol gradient fractionation of nuclear extracts (**Extended Data Fig. 2e-f**)^14^. The relative prevalence of GBAF, CBAF, and PBAF in nuclear extracts must be interpreted with the understanding that these assays are performed on the populations of nuclear proteins that are not tightly bound to insoluble chromatin. Specifically, the preponderance of SS18::SSX-bearing GBAF that is avidly bound to insoluble chromatin leads to GBAF’s underrepresentation in the non-chromatin-bound nuclear proteins resolved by the glycerol gradient. Notably, the CBAF-specific subunit ARID1A is difficult to identify at all in SyS nuclear protein gradients, but is no more prevalent in the chromatin bound fraction, as we previously reported^14^.

GBAF peaks were observed to be broader than CBAF or PBAF peaks (p = 2.88×10^-13^, p = 0.035, 2-tailed heteroscedastic t-tests, respectively for GBAF-to-CBAF and GBAF-to-PBAF), especially at TSSs, but also comprised the great majority of distal peaks for all BAF complexes (**Fig. 2c**). Profiling of BAF components at SAT and MAT pattern promoters identified striking contrasts, with CBAF and PBAF narrowly occupying the TSS of both types of promoters, and stronger fusion and GBAF enrichment at SAT pattern than at MAT pattern (H2AK119ub-lacking) or SST promoters (**Fig. 2d-f**, **Extended Data Fig. 2g-h**). Distal loci with H2AK119ub-called peaks show strong enrichment for H2AK119ub, fusion and GBAF. However, even loci lacking called peaks for H2AK119ub demonstrated the presence of GBAF and fusion in a similar, albeit diminished enrichment pattern (**Extended Data Fig. 2i-l**). These data suggest that SS18::SSX distributes with GBAF primarily to the regulatory elements of sarcomagenesis-associated genes. Notably, CBAF experienced a near-total depletion at distal loci, positions where it typically plays a vital role in remodeling chromatin at enhancers across various cell types, including stem cells (**Fig. 2g**)^25^. The narrow pattern of CBAF at the TSS (often with accompanying fusion), correlates with gaps in the H2AK119ub and H3K4me3 distribution; GBAF instead co-distributes with H2AK119ub in promoter CpG islands, identified loosely by high GC content of these promoter sequences (**Extended Data Fig. 2m-n**). Hypomethylated CpG islands identify developmental genes for bivalency as we report in our related study in human SyS^16^. In this fashion, H2AK119ub enrichment inversely correlated with CBAF presence among fusion peaks (**Extended Data Fig. 2o**).

### Synovial sarcoma transcriptional trajectories define gene sets

In order to profile the transcriptional changes that accompany SyS development along disparate epithelial, mesenchymal, or poorly differentiated trajectories, we performed single cell transcriptome profiling (scRNA-seq) on mouse SyS tumors comparable to those profiled epigenomically (**Extended Data Fig. 1a-b**). The tumors exhibited a variety of histomorphologies, indicated in hematoxylin and eosin tumor sections of tissue immediately adjacent to that submitted for scRNA-seq (**Fig. 3a**, **Extended Data Fig. 3a**). Clusters of tumor-infiltrating fibroblasts, endothelial cells, and immune cells were easily identified by their expression signatures (**Fig. 3b-c**, **Extended Data Fig. 3b**). Spatial profiling of transcription in additional mouse SyS tumor samples identified reference genes across histologically-specified cell types and permitted assignment of clustered malignant cells to specific categories (**Fig. 3d-f**, **Extended Data Fig. 4**). We then performed a pseudo-time trajectory from the likely earliest transformed cells to those that displayed mesenchymal spindle cell morphology, epithelial cell morphology, and poorly differentiated morphology (**Fig. 3g**). These trajectories revealed positively associated genes in both the SAT pattern (sarcomagenesis linked) and MAT pattern (housekeeping) gene sets (**Fig. 3h**, **Extended Data Fig. 5**).

**Fig. 3.**
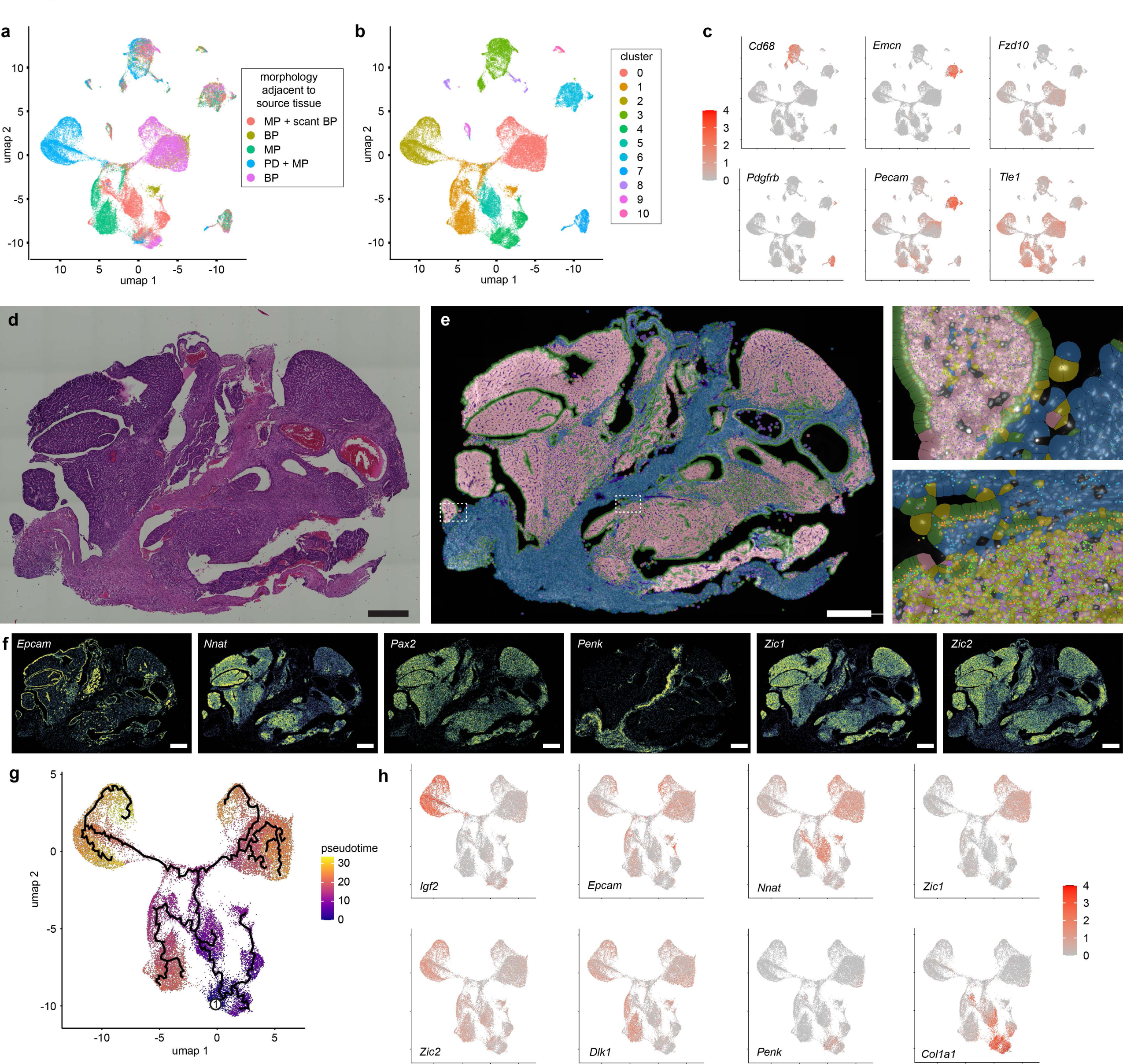
Single cell transcriptomics reveal gene sets associated with specific differentiation states of synovial sarcoma cells. (a) Seurat clustering by umaps in 2 dimensions noting 5 source samples (MP, monophasic; BP, biphasic; PD, poorly differentiated) and clusters (b) of mouse synovial sarcomas initiated by SS18::SSX2 expression by TATCre limb injection at 8 days of life. (c) Heatmaps for expression of marker genes across the clusters, identifying cluster 3 as monocytes/macrophages, 7 as fibroblasts, 6 as endothelial cells, and 0, 1, 2, 4, 5 as SyS cells, 8, 9, and 10 represent other supporting non-neoplastic cell types. (d) H&E stained photomicrograph of Xenium spatial transcriptomics sample of an hSS2 mouse tumor. (e) Xenium cluster calls for SyS cell types of monophasic (blue), epithelial and poorly differentiated (pink), glandular epithelial cells (green), endothelial cells (purple) with expanded sections demonstrating the positional expression of *Penk* (cyan sunbursts), *Epcam* (orange sunbursts), *Nnat* (purple squares), and *Zic1* (green Xs) transcripts. (f) Xenium heatmap expression data for marker genes of distinct histological cell types. (g) Whole transcriptome scRNA-seq of neoplastic cells clusters demonstrate divergent pseudotime trajectories into the varied Seurat umap clusters, beginning at the circled 1. (h) Heatmap expression of individual marker genes that distinguish between different neoplastic cell clusters.

### *Smarcb1* disruption disables the expression of SS18::SSX-driven developmental target genes

Next, we sought to determine the functional contribution of the narrow CBAF and PBAF localization at TSSs to SyS tumor phenotypes. Both of these BAF sub-families incorporate SMARCB1 and ARID components. We therefore began by interrogating the impact of *Smarcb1* genetic disruption^26^ in *hSS2*-driven tumors (**Extended Data Fig. 6a-b**). We previously demonstrated that *Smarcb1* knockout significantly shortened latency to tumorigenesis and altered cell morphology away from typical SyS features, with a distinct increase in nuclear atypia (**Fig. 4a-b**, **Extended Data Fig. 6c-d**)^14^. This suggests that biology associated with SMARCB1 loss is not included in fusion-only-driven oncogenesis. Transcriptional profiling by RNA-seq demonstrated significantly reduced expression of the majority of SAT pattern genes, inconsistent directional change in MAT pattern genes, and a subtle further reduction in some of the SST genes, which are already relatively lowly expressed in the *Smarcb1* wildtype tumors (**Fig. 4c-d**, **Extended Data Fig. 6e-i**). We confirmed that the loss of SMARCB1 leads to profound loss of PBAF complexes and a restored preponderance of CBAF complexes (**Fig. 4e**, **Extended Data Fig. 6j-k**), further confirming our prior observation that CBAF complexes can readily assemble in the absence of the SMARCB1 subunit^14^. We previously interpreted similar findings as demonstrating that PBAF loss at the TSS was solely responsible for the acute alteration in tumor phenotype. However, considering the tight correspondence of CBAF and PBAF distributions across the SyS genome, we further propose that CBAF fails to compensate PBAF loss at the TSS of SAT pattern gene promoters in the context of SMARCB1 loss. A conspicuous dip at these sites in *Smarcb1*-silenced tumors represents focal loss of general BAF complex ChIP-seq signal here. This dip strongly mirrors that of GBAF distribution in *Smarcb1*-wildtype tumors (**Fig. 4f**, **Extended Data Fig. 6l-o**). Taken together, our data suggest that focal absence of GBAF binding near the TSS in SyS tumors is compensated by general BAF binding of TSS-localized CBAF and PBAF, both of which are lost in *Smarcb1*-silenced tumors.

**Fig. 4.**
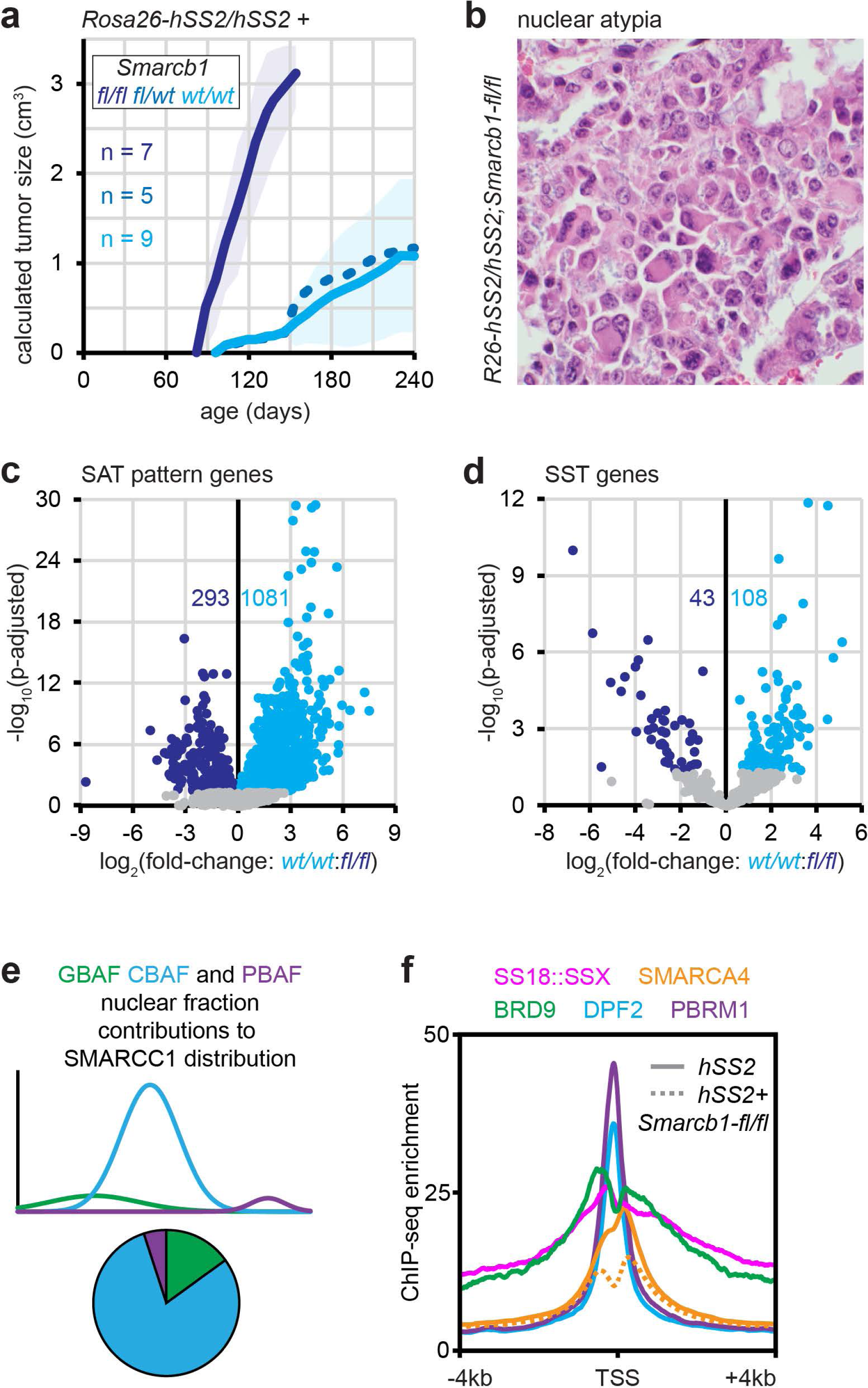
SMARCB1 loss limits PBAF assembly in SyS and blocks CBAF from TSS binding. (a) Growth trajectories of mean tumor size (shading indicates ± standard deviations, p-values from two-tailed, heteroscedastic *t* tests) in *hSS2* mice injected with TATCre at day 28 of life, comparing littermates with varied *Smarcb1-fl* genotypes. (b) Example photomicrograph of the bizarre nuclear atypia apparent in an *hSS2* tumor with homozygous floxed *Smarcb1*. (c) Differential expression (by bulk tumor RNA-seq) of SAT pattern genes and (d) SST pattern genes of *hSS2* tumors with *wildtype* versus homozygous floxed *Smarcb1*. (e) Graphical histograms indicating the relative contribution of GBAF, CBAF and PBAF in glycerol nuclear fractions of an *hSS2*, *Smarcb1-*disrupted tumor. (f) ChIP-seq enrichment plots for BAF components in *hSS2*, *Smarcb1* wildtype tumors at promoters defined by the focal loss of SMARCA4 enrichment at the TSS in combination genotype tumors.

### *Pbrm1* disruption blocks SS18::SSX reprogramming, encouraging alternate oncogenic pathways

Silenced expression of a PBAF specific subunit, *Pbrm1*^27^ in *hSS2*-driven tumorigenesis also led to faster tumor development (**Fig. 5a**, **Extended Data Fig. 7a-c**). *Pbrm1*-silenced tumors lacked the full array of SyS histomorphologies observed in *Pbrm1*-wildtype tumors, instead exhibiting increased nuclear atypia (**Fig. 5b**, **Extended Data Fig. 7d**). Compared to the negative impact of *Smarcb1* silencing on SAT pattern gene expression in SyS tumors, *Pbrm1* silencing was comparatively subtle (**Fig. 5c**). This argues that CBAF (with SMARCB1 intact) can compensate for PBAF disruption at the narrow TSS binding positions in SyS, remodeling chromatin to enable gene expression. More striking was the reduced expression of MAT pattern genes in *Pbrm1*-silenced tumors (compared to *Smarcb1* silencing) (**Fig. 5d**). *Pbrm1* silencing in these *hSS2* tumors achieved the anticipated reduction in the presence of PBAF complexes in nuclear extracts and the loss of *Pbrm1* transcripts by RNA-seq (**Fig. 5d-e**, **Extended Data Fig. 7e-f**).

**Fig. 5.**
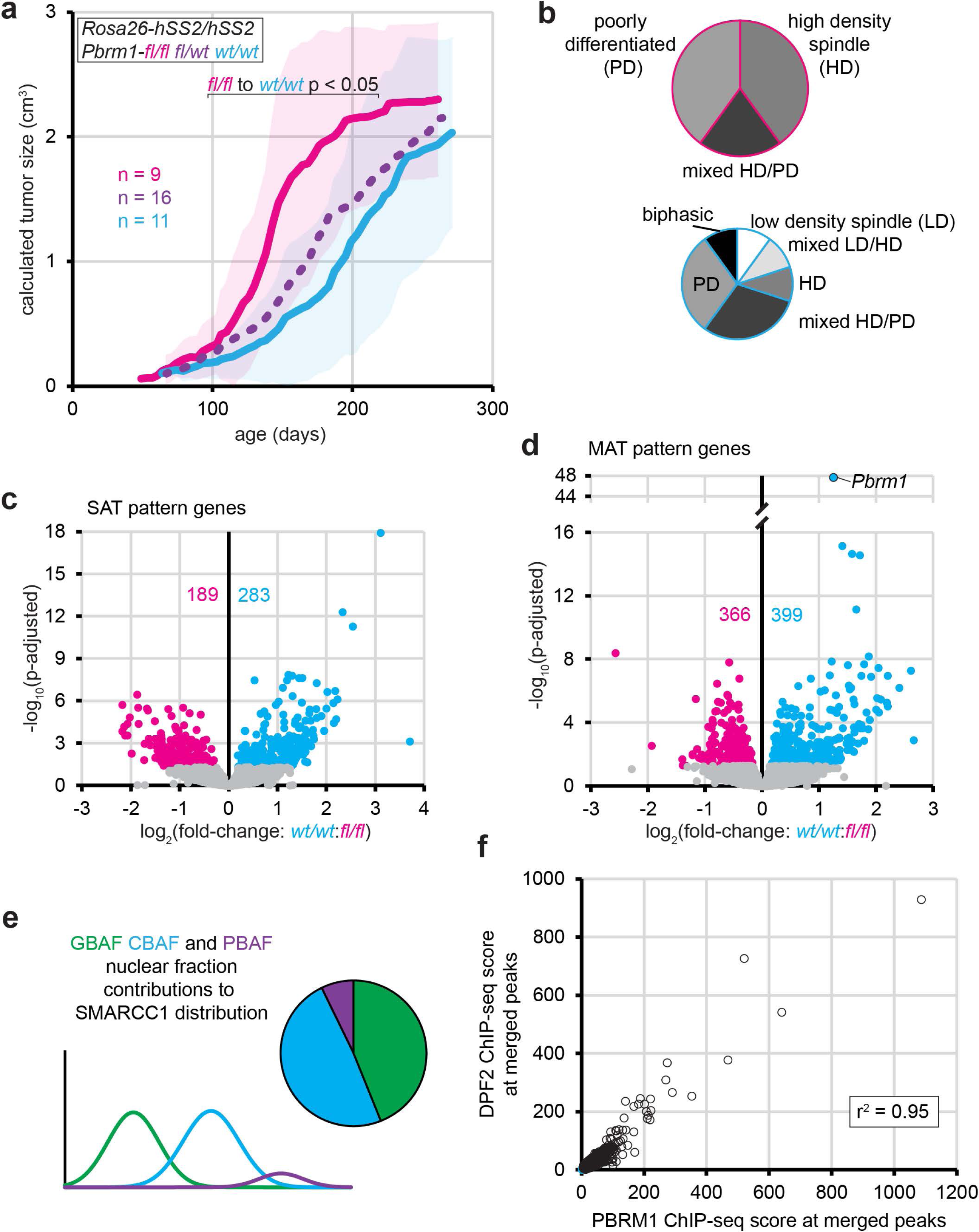
PBRM1 loss prohibits PBAF assembly and blocks the synovial sarcoma phenotype. (a) Growth trajectories of mean tumor size (shading indicates ± standard deviations, p-values from two-tailed, heteroscedastic *t* tests) in *hSS2* mice injected with TATCre at day 8 of life comparing littermates with varied *Pbrm1-fl* genotypes. (b) Pie charts indicating the histomorphology of *hSS2*;*Pbrm1-fl/fl* versus *hSS2*;*Pbrm1-wt/wt* tumors (n = 15, n = 10). (c) Differential expression (by bulk tumor RNA-seq) of SAT pattern genes and (d) MAT pattern genes of *hSS2* tumors with *wildtype* versus homozygous floxed *Pbrm1*. (e) Graphical histograms indicating the relative contribution of GBAF, CBAF, and PBAF to SMARCC1 distributions in glycerol nuclear fractions of an *hSS2*, *Pbrm1*-disrupted tumor. (f) DPF2 versus PBRM1 ChIP-seq enrichment across merged peaks between the two.

Although CBAF and PBAF ChIP-seq enrichments are strongly correlated across the entire genome (**Fig. 5f**), the discrepancy between the numbers of MAT genes (more) versus SAT genes (fewer) showing decreased gene expression, suggests that CBAF (with SMARCB1 intact) is more capable of compensating for PBAF absence at developmental TSSs than at housekeeping genes.

For optimal rigor, we also performed the experiments in the *hSS1* background (expressing SS18::SSX1 from *Rosa26* via TATCre injection, which drives slightly slower synovial sarcomagenesis.)^13^ While *Pbrm1* loss decreased latency to tumorigenesis in *hSS2* tumors, homozygosity for floxed *Pbrm1* in *hSS1* tumors paradoxically slowed tumorigenesis (**Extended Data Figure 7g-j**). Tumors arising more slowly in *hSS1;Pbrm1-fl/fl* mice retained normal SyS histomorphologies and lacked the nuclear atypia seen in *hSS2;Pbrm1-fl/fl* (**Extended Data Fig. 7k**). Tumors developing in *hSS1;Pbrm1-fl/fl* mice also depleted *Pbrm1* expression (by RNA-seq) less than did *hSS2* tumors (**Extended Data Fig. 7e,l**), suggesting that tumor cells recombining only one of the two floxed *Pbrm1* alleles experienced stronger positive selection in the *hSS1* background. Principal component analysis (PCA) of entire transcriptomes determined by RNA-seq of *Pbrm1-fl/fl* and *wt/wt* tumors from both *hSS1* and *hSS2* mice demonstrated disparate clustering of the *hSS2;Pbrm1-fl/fl* tumors, but retained clustering of *hSS1;Pbrm1-fl/fl* tumors with *Pbrm1* wildtype tumors from both backgrounds (**Extended Data Fig. 7m**). Overall, this apparent paradox suggests that *Pbrm1* silencing similarly blunted synovial sarcomagenesis programs in both SSX::SS18 backgrounds. This blunting of SyS programs tipped the faster growing *hSS2* tumors^13^ into an alternate tumorigenesis program, but merely slowed *hSS1* tumor development, causing selection for alleles that escaped recombination.

### *Arid1a* or *Arid1b* disruption enhances the transcriptional silencing derived from SS18::SSX expression

Silencing of *Arid1a* or *Arid1b* sped tumorigenesis in both backgrounds (insignificantly in *hSS2*/*Arid1b*-loss mice, but significantly in all others), without significant change in the histomorphology of the developing tumors (**Fig. 6a-e**, **Extended Data Fig. 8a-p**)^28,29^. Tumors with homozygous disruption of *Arid1a*, especially, demonstrated rounder nuclei and higher cellular density (**Fig. 6e**). *Arid1a* or *Arid1b* silenced tumors still exhibited strong epithelial features, with or without epithelial gland formation (**Fig. 6e**, **Extended Data Fig. 8d,h,l,p**).

**Fig. 6.**
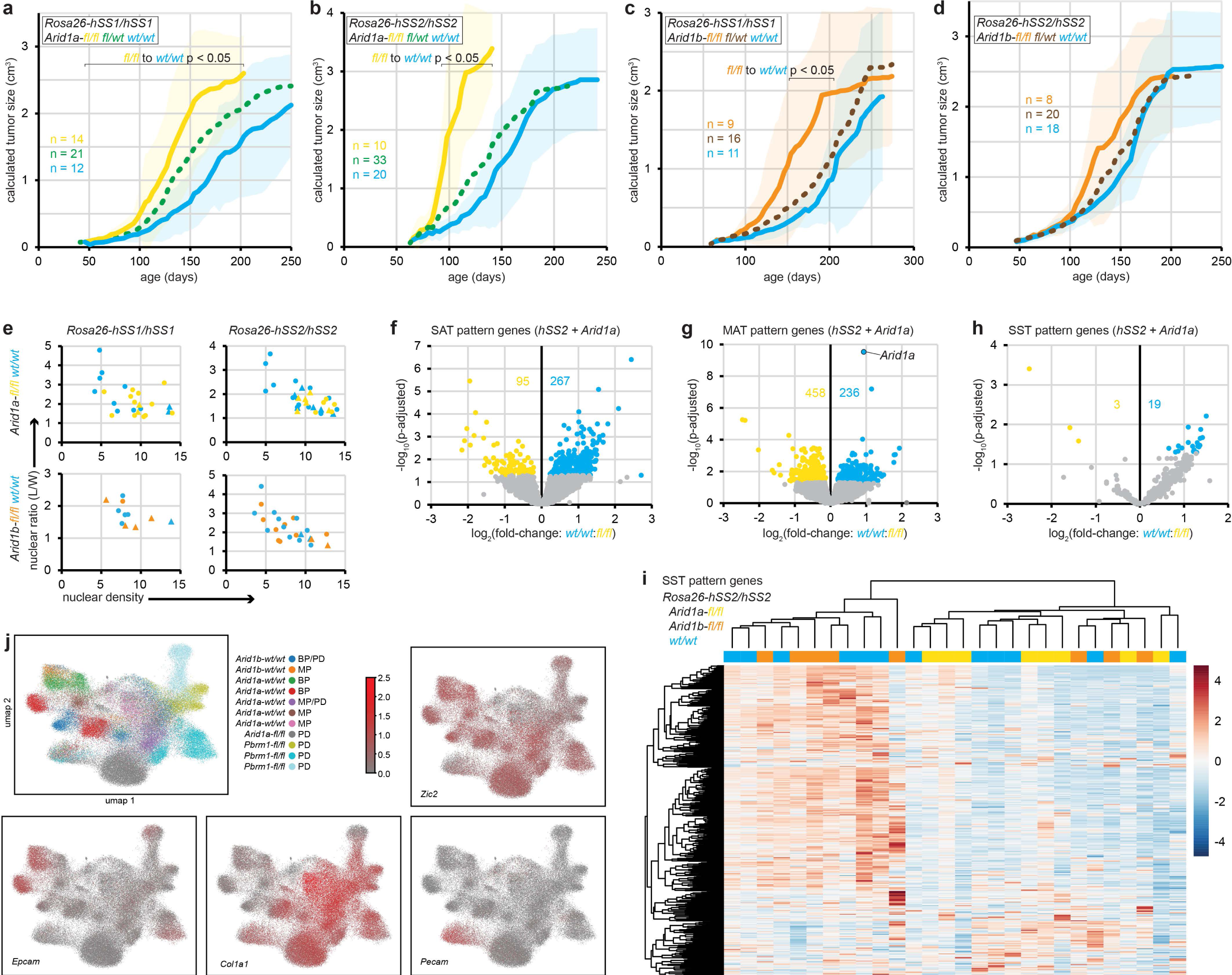
CBAF disruption enhances tumorigenesis without compromising the synovial sarcoma phenotype. (a) Growth trajectories of mean tumor size (shading indicates ± standard deviations, p-values from two-tailed, heteroscedastic *t* tests) in *hSS1* and (b) *hSS2* mice injected with TATCre at day 8 of life comparing littermates with varied *Arid1a-fl* genotypes. (c) Similar tumor growth curves for *hSS1* and (d) *hSS2* for littermates with varied *Arid1b-fl* genotypes. (e) plots of nuclear shape ratios (closer to 1 indicates roundness) and density for tumors from the different genetic cohorts, with monophasic and poorly differentiated tumors across the spectrum in round dots and tumors with epithelial histomorphology indicated by triangles. (f) Differential expression of SAT, (g) MAT, and (h) SST pattern genes comparing *hSS2* tumors with *wildtype* versus homozygous floxed *Arid1a* genotypes. (i) expression heatmap of bulk RNA-seq for SST pattern genes among *hSS2* tumors with either *Arid1a* homozygous floxed, *Arid1b* homozygous floxed, or *wildtype* for both. (j) Xenium In Situ single cell transcriptomic umap plots indicating specific clusters that are epithelial or mesenchymal neoplastic populations of cells, versus endothelial cells, and how much each of the unique sample sources contribute to the varied clusters.

Although there was no appreciable further depletion of CBAF complexes in *Arid1a*-silenced tumors beyond that of wildtype *hSS2* tumors, there was also no apparent compensatory increase in expression of either paralog upon deletion of the other (**Extended Data Fig. 9a-b**). Similar to *Pbrm1*-silenced *hSS2* tumors, more of the SAT pattern genes had higher expression in *Arid1a*-wildtype *hSS2* tumors than in tumors with silenced *Arid1a*, but the difference was even more subtle (**Fig. 6f**). MAT genes were similarly not impacted to any large degree (**Fig. 6g**), unlike *Pbrm1*-silenced *hSS2* tumors in which MAT genes were lost. The most striking pattern of altered transcription was in the SST genes, with *Arid1a* disruption reducing expression further, even in this gene set defined by silenced transcription during synovial sarcomagenesis (**Fig. 6h-i**). Moreover, among the genes exhibiting increased expression along the pseudotime trajectory, most of those that were expressed at higher levels in *Arid1a*-silenced tumors had MAT pattern promoters (**Extended Data Fig. 9c**). Overall, *Arid1a* or *Arid1b* disruption rendered tumor transcriptomes that clustered within the *hSS2* tumor transcriptomes, unlike *Pbrm1* disrupted tumors (**Extended Data Fig. 9d**). Similarly, Xenium assessments of *Pbrm1*-silenced tumors showed a wider departure from the core clusters of neoplastic cell types than an *Arid1a*-silenced tumor (**Fig. 6j**, **Extended Data Fig. 10**)

## Discussion

Like SS18, the SS18::SSX fusion incorporates into both GBAF and CBAF (not PBAF), and is found coincident with both subfamilies of BAF complexes on chromatin, but has distinct relationships at gene regulatory elements bearing specific epigenomic patterns with each of these complexes.

Fusion oncoprotein distribution follows H2AK119ub patterns, displaying broad peaks both within and outside promoters, and appears to be the driving force behind GBAF distribution (**Fig. 7**).

**Fig. 7.**
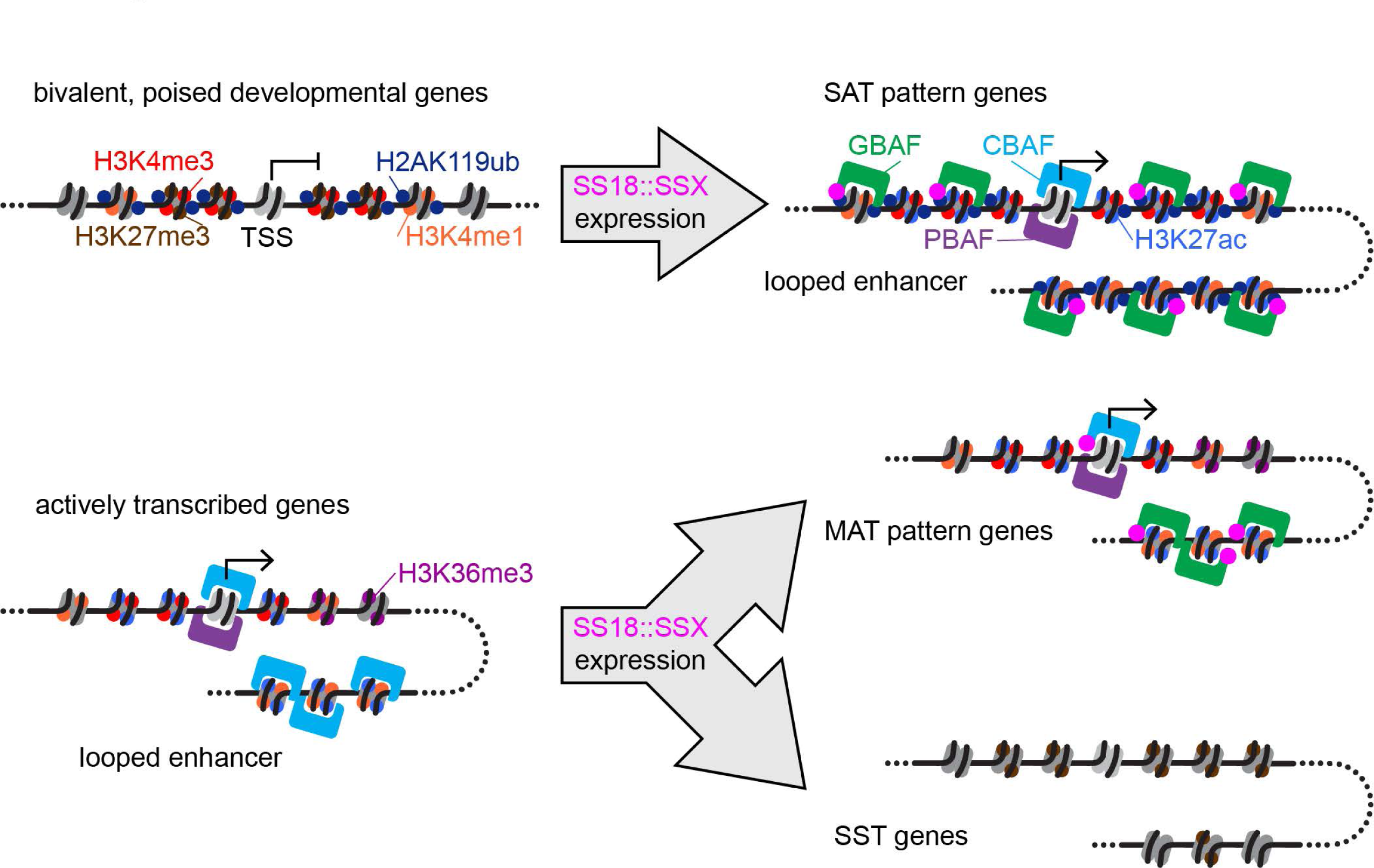
Working model of transcriptional regulation by SS18::SSX. Genes with promoters and enhancers that are poised, exhibiting H3K27me3/H3K4me3 bivalency in development, are shifted towards expression by the presence of SS18::SSX in cells, which promotes H3K4me3 monovalency following the redistribution of GBAF complexes containing the fusion. Other genes that are expressed at baseline, are silenced during sarcomagenesis from expression of the fusion and subsequent CBAF desertion from its typical distribution across the genome-wide chromatin of normal cells. Other genes maintain expression and retain CBAF enrichment, but only narrowly near TSSs, where PBAF is also present.

Following fusion expression, this gain of targeting and functional activity for GBAF complexes involves promoters (and distal elements) that transition from bivalency in progenitors to active histone marks in the tumor cells^15^, likely in part through chromatin opening by GBAF. The changing patterns of histone marks over time and tumor development were demonstrated through comparisons with a known cell of origin in our other mouse experiments^15^. The particular patterns identified across a series of developmental target genes supports interpretations of the human profiles of SyS tumors that exhibit variable retention of bivalency for H3K27me3 and H3K4me3 versus complete conversion to H3K4me3 coincident with H2AK119ub, though curiously lacking H3K27me3^16^. Here, we demonstrate that the strong relationship between H2AK119ub and H3K4me3 is at least coincident—if not driven by—the presence of GBAF (with incorporated SS18::SSX). These genes, that we designated as SAT pattern genes, comprise the direct target genes of the fusion oncoprotein, which are driven toward elevated transcription. Gene targets of GBAF+fusion comprise an activated expression signature that is strongly shared across species (human and mouse)^15,16^. As much as these SAT pattern genes are targeted by GBAF+fusion, their expression also depends on the functional presence of either PBAF or CBAF (each incorporating SMARCB1) narrowly at the TSS. Whereas GBAF is apparently retained at SAT promoters when *Smarcb1* is silenced, this silencing drives promoter desertion by CBAF and PBAF, coincident with transcriptional attenuation. This finding suggests that GBAF+fusion has limited capacity to bind to and remodel chromatin at the TSS itself, and instead may open the region for the action of transcription factors, CBAF and PBAF. Therefore, SAT target genes that are transcriptionally activated by the fusion oncoprotein itself are marked by H2AK119ub, H3K4me3, and GBAF, but additionally require PBAF or CBAF at the TSS.

While the fusion appears to drive the distribution of GBAF across the genome as just outlined, the distribution of CBAF+fusion complexes appears to be directed primarily by CBAF itself, with fusion accompanying as passenger. Consistently, CBAF+fusion is found in narrow peaks near TSSs of promoters lacking H2AK119ub altogether, or at least lacking H2AK119ub focally near TSSs. These are not normal CBAF targets, as the typically distal enhancer positions of CBAF are almost completely abrogated in SyS. There are many potential explanations for this that will be pursued in further experiments.

The overall reduction in CBAF complex protein levels directed by SS18::SSX is importantly incomplete^14^, as a small proportion of CBAF complexes remains, usually bearing the fusion oncoprotein SS18::SSX and SMARCB1. Importantly, we report the near complete redistribution of these CBAF complexes on chromatin. The observation that disruption of ARID1A, a critical subunit of CBAF, decreased latency to tumorigenesis without significantly altering tumor phenotype argues that CBAF-depletion only furthers or enhances aspects of the tumorigenesis process that are otherwise intrinsic to SS18::SSX expression. While one might suggest that ARID1B simply compensates for ARID1A absence in these cells, blunting the effect of ARID1A loss, the loss of ARID1B had an even smaller effect on tumors. There was no noticeable change in the expression of either component when the other was silenced.

Taken together, our work reveals two main and distinct mechanisms for SS18::SSX-mediated transcriptional reprogramming via BAF complexes in SyS: 1) upregulation of gene targets of GBAF with SS18::SSX, and 2) silencing/attenuation of CBAF targets due to diminished and redistributed CBAF. Here, transcriptional silencing likely involves loss of typical CBAF distributions to distal regulatory elements, which is enhanced by *Arid1a* loss. However, we note the presence of additional genes, whose transcription is upregulated by this disrupted CBAF mechanism. One possible group of such genes would be the MAT pattern genes which were found to associate with pseudotime trajectories mapped onto the development of more fully reprogrammed cells. While half of the pseudotime trajectory genes were SAT pattern genes, the other half were MAT pattern, or housekeeping, genes and were largely upregulated by genetic disruption of *Arid1a*. The effect of CBAF dysregulation within cells expressing the fusion may vary across different cells of origin and possibly contribute heavily to the cell death phenotype driven by SS18::SSX expression in many cell types. This may be attributable to the toxicity associated with BAF disruptions, and raises the interesting question regarding how SyS progenitors in vivo can survive and adapt to SS18::SSX fusion expression.

Notably, any additional genetic perturbation added to SS18::SSX expression led to at least slight reductions in the expression of SAT pattern genes. Although profoundly reduced expression of these genes in tumors lacking *Smarcb1* may be explained by loss of PBAF and CBAF at the TSS (where GBAF+fusion is possibly incapable of binding), the disruptions of either PBAF or CBAF by other subunit silencing experiments also diminished expression of subsets of SAT pattern genes. This observation fits a model for heterogeneity in human SyS, where tumors that have more copy number variation also have more retained bivalency, as shown in our human datasets^16^. It may be that CBAF loss (or alternative oncogenic mechanisms) in SyS reduce the dependence on GBAF+fusion for complete reprogramming at developmental genes, since oncotransformation may also be fundamentally enabled by CBAF-induced transcriptional dysregulation.

## Methods

### Animal Studies

Tumor growth rates in mice carrying *Arid1a-fl/fl*^29^, *Arid1b-fl/fl*^28^, or *Pbrm1-fl/fl*^27^, on an *hSS2* (*SS18::SSX2*)^12^ or *hSS1* (*SS18::SSX1*)^13^ background were interrogated. Spontaneous tumors with each combination genotype were generated by limb injection with TATCre protein at 8 days of age. Caliper measurements started when tumors became visible. Tumor volumes were calculated using the following formula: tumor volume = (D x d^2^)/2, in which D and d refer to the long and short tumor diameter, respectively. All males and females from experimental litters were included in each group. Genotyping primer sequences are in the Resources Table.

### Histology

Time of morbidity was variable across individuals and groups, at which point mice were euthanized and tumors harvested at necropsy. Tissues were fixed in 4 percent paraformaldehyde, dehydrated in increasing ethanol gradients, embedded in paraffin, and sectioned at 10μm thickness for standard H&E staining. For the quantitative assessments of histological features in the different groups, slides were reviewed after randomization for order, blinded regarding the genotype of the mouse from which the tumor was harvested. Nuclear ratios were determined dividing the length of the long axis of each nucleus over the length of the orthogonal axis. These were averaged from three measurements in each histomorphologically distinct area of the tumor section, then multiplied by the fractional contribution of that histomorphological area to the whole section. Nuclear density was calculated for each histomorphologically distinct region as a count of nuclei in a 500 µm^2^ square on the photomicrograph.

### Mouse tumor nuclear extraction

Nuclear extraction was performed according to our previously published protocols^14^. Briefly, tumors were pulverized at −80°C in Covaris TT1 tissue tubes (Covaris). Tissue pellets were washed and lysed with Buffer A (20 mM HEPES pH 8.0, 1.5 mM MgCl2, 10 mM KCl, 0.25% NP-40, 0.5 mM DTT with 2x protease inhibitor cocktail). Pelleted nuclei were then resuspended in Buffer C (20 mM HEPES pH 8.0, 25% glycerol, 1.5 mM MgCl2, 420 mM KCl, 0.25% NP-40, 0.2 mM EDTA, 0.5 mM DTT, and 2x protease inhibitor cocktail) and disrupted using a dounce homogenizer (Sigma).

### Density sedimentation gradients and calculation of BAFs abundances

Density sedimentation gradients were performed according to our previously published protocol with some minor changes^14^. Nuclear extractions from tumors were diluted 1:1 with dilution buffer (20 mM HEPES pH 8.0, 1.5 mM MgCl2, 0.2 mM EDTA, 0.5 mM DTT, and 2x protease inhibitor cocktail). Samples were loaded onto tubes (Beckman) with 11 ml 10-30% glycerol gradients containing 20 mM HEPES pH 8.0, 1.5 mM MgCl2, 200 mM NaCl, 0.2 mM EDTA, 0.5 mM DTT, and 2x protease inhibitor cocktail. Tubes were then loaded into a SW41 rotor and centrifuged at 40,000 rpm for 20 hours at 4°C. 24 fractions were automatically collected with a BioComp fractionation system (BioComp), with each fraction containing 466 µL of sample. Fractions were run on an SDS-PAGE gel for Western blot analysis. The abundances of different BAF-family subtypes were calculated using our previously published protocol^14^.

### Western blotting

Primary antibodies used for western blots are listed in the Resources Table. Tumor protein samples were separated by 10% PAGE gel electrophoresis and transferred to PVDF membranes. Membranes were blocked with 5% milk TBST and incubated with primary antibodies overnight. After four TBST washes, blots were incubated with HRP-conjugated species-specific secondary antibodies, washed four times with TBST and developed with SuperSignal^TM^ West Dura Extended Duration Substrate (Thermo).

### RNAseq

Fresh frozen tumor tissue was disrupted using a Tissue-Tearor (BioSpec) in 1ml of TRIzol™ reagent (ThermoFisher). 200 µL isopropanol was added to the sample, followed by brief vortexing, and centrifugation. Supernatants were transferred into fresh tubes, to which 1 volume of Ethanol was added. Further RNA cleanup was performed using Direct-zol^TM^ RNA Miniprep kit (Zymo), starting at step 2 of the Zymo protocol. RNA-seq libraries were prepared with the NEBNext Ultra II Directional RNA Library Prep with rRNA Depletion Kit, and sequenced on a NovaSeq using the 2 x 150 bp protocol (Illumina) for approximately 25-30 million reads per sample. Read alignment was accomplished with the STAR (2.7.11) against the mm10 version of the mouse genome^30^. Count matrices were generated using featureCounts (version 1.6.3), while differential expression analysis was performed with DESeq2 version 3.11^31^. Genes displaying a false discovery rate (FDR) below 0.05 were considered statistically significant. For the purpose of visual representation, expression levels were transformed to regularized logarithm (rlog) counts. Principal Component Analysis (PCA) plots were derived from the first two principal components based on the rlog-transformed data of the selected gene set. Additionally, heatmaps reflecting this gene set were created through unsupervised hierarchical clustering, utilizing sample Euclidean distances.

### Chromatin Immunoprecipitation (ChIP)

ChIP-seq was performed according to our previously published protocols^14^. Snap frozen tumor specimens were pulverized as previously described. Tumor pellets were crosslinked with 1% formaldehyde and quenched with 0.125 M of glycine. For double cross-linking (on samples undergoing ChIP with DPF2, PBRM1, and control samples of for SS18::SSX and ARID1A), pulverized tumors were thawed in HBSS and cross-linked with fresh 20 mM dimethyl pimelimidate (dissolved in 0.2 M Triethanolamine pH 8.2) prior to formaldehyde crosslinking.

After three washes in cold PBS, nuclei were incubated in mild lysis buffer (10 mM Tris-HCl pH8.5, 10 mM NaCl, 0.5% NP-40) containing protease inhibitor (PI, Sigma), followed by wash buffer (10 mmol/L Tris-HCl pH8.5, 200 mM NaCl, 1mM EDTA and 1% SDS with PI), and strong lysis solution step (50 mM Tris-HCl pH8.0, 10mM EDTA and 1% SDS with PI) immediately followed by dilution 1:10 with buffer (16.7 mM Tris-HCI pH8.1, 16.7 mM NaCl, 1.2 mM EDTA, 1.1% Triton X-100, 0.01% SDS with PI). Samples were then sonicated with an EpiShear Probe Sonicator (Active Motif). 5 µg of primary antibody was then added to the sonicated chromatin for incubation at 4°C overnight. The next day, 100 µL of washed Dynabead slurry was incubated with the IPed chromatin for 4.5 hours. Following subsequent wash and reverse crosslinking steps, DNA was purified with a DNA clean & concentrator kit (Zymo). Sequencing libraries were prepared with the NEBNext Ultra II DNA Library Prep Kit, and sequenced with a NovaSeq sequencing system using the 2 x 150 bp protocol (Illumina) for approximately 25-30 million reads per sample.

Native ChIP-seq was performed according to previously published protocols^32^.

Reads were aligned to mm10 mouse genome version using Novoalign (Version 3.00) for paired-end reads. Peaks were called from each of the aligned bam files against input reads using MACS2 ^33^, (version 2.2.9) with the parameters: callpeak -B --SPMR --qvalue = 1e-3 --mfold 15 100. ChIP input was used as the background for MACS2. MACS was used to produce normalized bedgraphs, which were subsequently converted to bigWig files. Peaks were filtered to remove peaks that are in blacklist, including ENCODE blacklisted regions ^34^. Duplicate reads were removed using samtools rmdup for all downstream analyses ^35^. Merged bigWig enrichment files for each condition with multiple replicas were generated in an average manner followed by normalization of read depth.

Heatmaps and profile plots that illustrate the scores corresponding to genomic regions were produced using the plotHeatmap function in deepTools (version 3.5.6) ^36^. This followed the computation of scores for each genomic region, a process which was executed using the computeMatrix function.

Correlation analyses of various ChIP-Seq datasets were conducted using the multiBigwigSummary function from the deepTools suite, focusing specifically on Brg1 binding sites. The Pearson correlation coefficient was employed to quantify the strength and direction of the relationship between the datasets.

### HiChIP

HiChIP assays were performed based on published protocols^14^ with several modifications. *hSS2* mouse tumors were harvested and digested to single cell suspension with the Miltenyi mouse tumor dissociation kit (Miltenyi Biotec). Approximately 5-10 million cells were crosslinked with 1% of formaldehyde, quenched with 0.125 M of glycine, and lysed with Hi-C lysis buffer (10 mmol/L Tris-HCl pH 7.5, 10 mmol/L NaCl, 0.2% NP40) supplemented with protease inhibitors.

Cross-linked chromatin was digested by MboI restriction enzyme. The digested overhangs were then filled with dCTP, dGTP, dTTP, and biotin-labeled dATP, followed by ligation with T4 DNA ligase. Nuclei were then resuspended in nuclear lysis buffer (50 mmol/L Tris-HCl pH7.5, 10 mmol/L EDTA, 1% SDS) supplemented with protease inhibitors and sonicated with Qsonica (Q800). The sonicated chromatin was then diluted with ChIP dilution buffer (16.7 mM Tris-HCl pH 7.5, 1.2 mM EDTA, 1.1% Triton X-100, 167 mM NaCl, 0.01% SDS) prior to DNA fragment capture with H3K27ac antibody. Streptavidin C1 magnetic beads were applied to capture ligated DNA fragments, and HiChIP libraries were prepared using Illumina Tagment DNA Enzyme and Buffer Kit. Sequencing was performed with the NovaSeq sequencing system using the 2 x 150 bp protocol (Illumina).

For alignment, the HiChIP sequencing data were mapped to the mouse mm10 reference genome utilizing the HiC-Pro pipeline ^37^. Chromatin loops were determined using the Hichipper pipeline ^38^ and those with an FDR value of less than 0.05 were selected as high-confidence loops for further investigation.

Promoter regions and distal loop anchors, each spanning a 4 kb length, were pooled to define the target genomic intervals. For every target, we calculated the cumulative score (which includes both fusion and H2AK119ub markers) of the loop-associated distal enhancers and promoter region.

### Single cell RNA-seq

*hSS2* mouse tumors were harvested at morbidity. Half of the tissue was digested (as described above) using the Miltenyi mouse tumor dissociation kit (Miltenyi Biotec). The remaining tissue from each sample was embedded in OCT for H&E staining in order to classify the corresponding tumor phenotype. After dissociation the cell suspension was filtered through a 40 µm strainer and treated with red blood cell lysis solution (Miltenyi Biotec). Lastly, cells were resuspended in PBS + 0.04% BSA at a concentration of ∼1,000 cells/μL for single-cell sequencing. 10,000 cells per sample were targeted. scRNA-seq libraries were prepared using Chromium Next GEM Single Cell 3ʹ Reagent Kits v3.1 according to the User Guide (CG000204), followed by amplification with paired-end dual-indexing (28 cycles Read 1, 10 cycles i7, 10 cycles i5, 90 cycles Read 2). The cellranger count command was employed for demultiplexing, barcode processing, and single-cell 3’ gene counting, resulting in a filtered matrix of gene expression counts for individual cells.

Quality control, normalization, and downstream analysis were performed using the Seurat package (v5.0.3) in R. Cells were filtered based on quality metrics, including removal of cells with mitochondrial features surpassing 25%, or total features below 750. Data were then normalized using the SCTransform function^39^. Cluster markers were identified using the FindMarkers function, with a minimum log-fold change of 1, pct.1 of 0.3 (expression of a gene in more than 30% of cells within the target cluster) and an adjusted p-value threshold of 0.01.

For pseudotime analysis, processed scRNA-seq data were imported into Monocle 3 (version 1.3.4) (http://cole-trapnell-lab.github.io/monocle-release/monocle3), and converted to cell data set format with the SeuratWrappers Package (version 0.3.2).

### Xenium In Situ

Seven *hSS2* mouse tumors (with histologically different morphology), one *hSS2; Ariad1a^fl/fl^* and three *hSS2; Pbrm1^fl/fl^* tumors were selected for Xenium In Situ analysis on FFPE tumor sections. 348 genes were designed for cell type identification (248 of Xenium pre-designed mouse Brain Panel and 100 customer designed SAT genes^15^). Five-μm-thickness FFPE sections were placed onto a Xenium slide, followed by deparaffinization and permeabilization for mRNA accessibility. Probe hybridization, Ligation and amplification were performed according to the Xenium In Situ GeneExpression User Guide (CG000582) prior to processing within the the Xenium Analyzer Instrumentation pipeline. Data generated by this pipeline was further analyzed downstream. Cell segmentation processing was performed using the cell segmentation user guide (CG000749), and post-Xenium tissue H&E staining was performed according to user guide CG000613.

Data integration and normalization steps were carried out in python package Scanpy ^40^ to correct for technical variance and to normalize expression counts. Spatial expression data were merged across different sections, with normalization to align gene expression levels.

### Single cell clustering and UMAP visualization

Cell clustering was performed using a shared nearest neighbor (SNN) modularity optimization algorithm ^41^. Louvain community detection was applied to the PCA-reduced data (30 principal components) to identify distinct transcriptional clusters within the tissue. For spatial data visualization, UMAP (Uniform Manifold Approximation and Projection) was applied to generate two-dimensional embeddings that reflect the spatial gene expression patterns.

### Statistical analysis

For the comparison of means between two independent groups, we utilized the two-tailed Student’s t-test, as facilitated by GraphPad Prism software (version 9.0). We established statistical significance at p-values of less than 0.05 or 0.01, which are specified accordingly in the legends accompanying each figure. All data are depicted as the mean ± standard deviation. The determination of sample sizes was guided by the variability observed in preliminary experiments. Notably, the numbers (n) referenced in the figure legends correspond to the count of biological replicates rather than repeated measures of the same specimen. We have employed additional statistical methods as necessary, with the specifics of these analyses duly noted within the legends of each figure. Furthermore, statistical procedures pertinent to genomic data have been detailed in the respective analysis subsections under the Methods section.

Non-parametric Kruskal-Wallis H tests were applied to assess the statistical significance of independent groups where the assumption of normality was not met. Dunn’s test was used for post-hoc analysis after a Kruskal-Wallis H test indicated a significant result. Bonferroni correction was used to adjust the significance levels.

## Resources Table

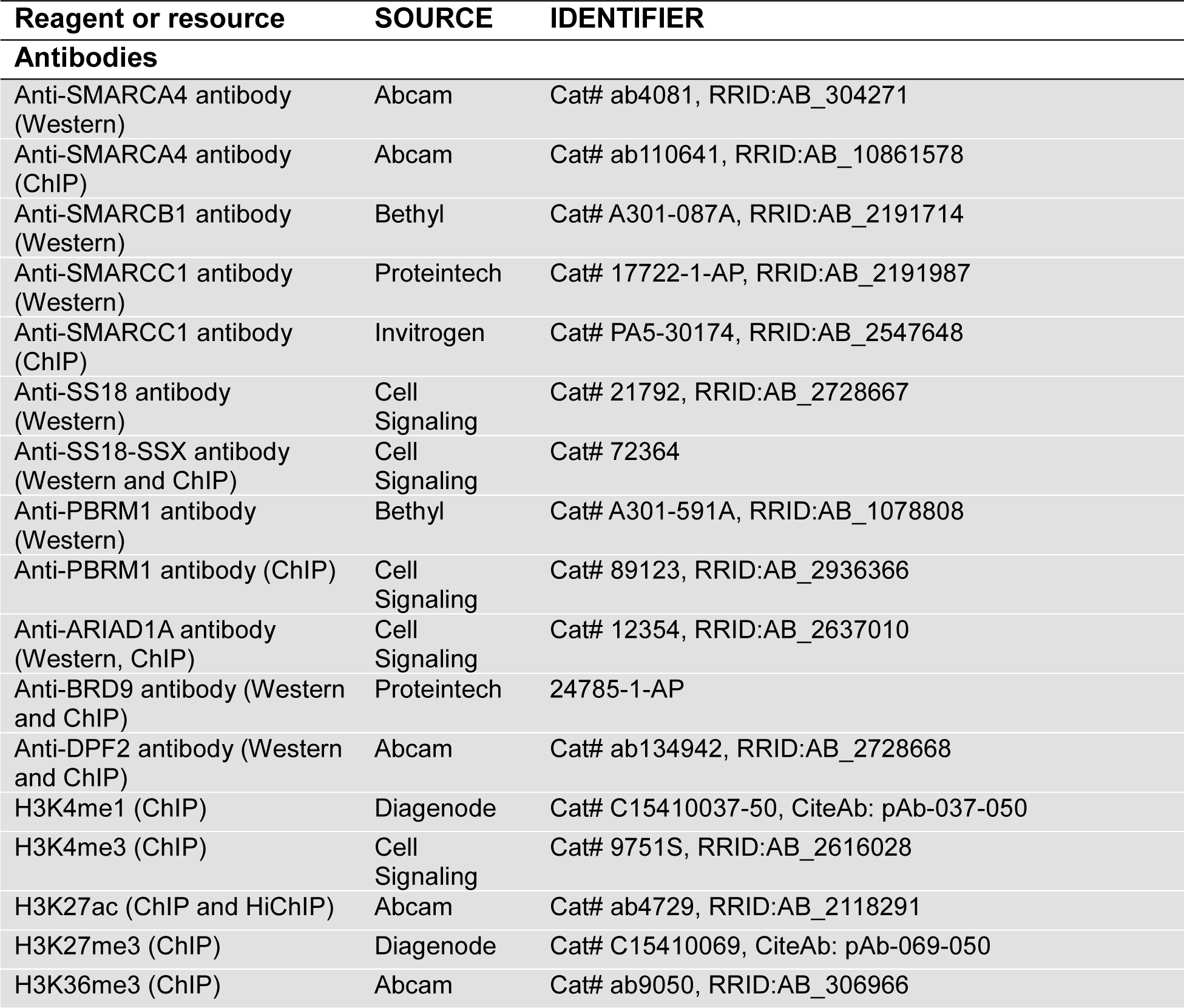
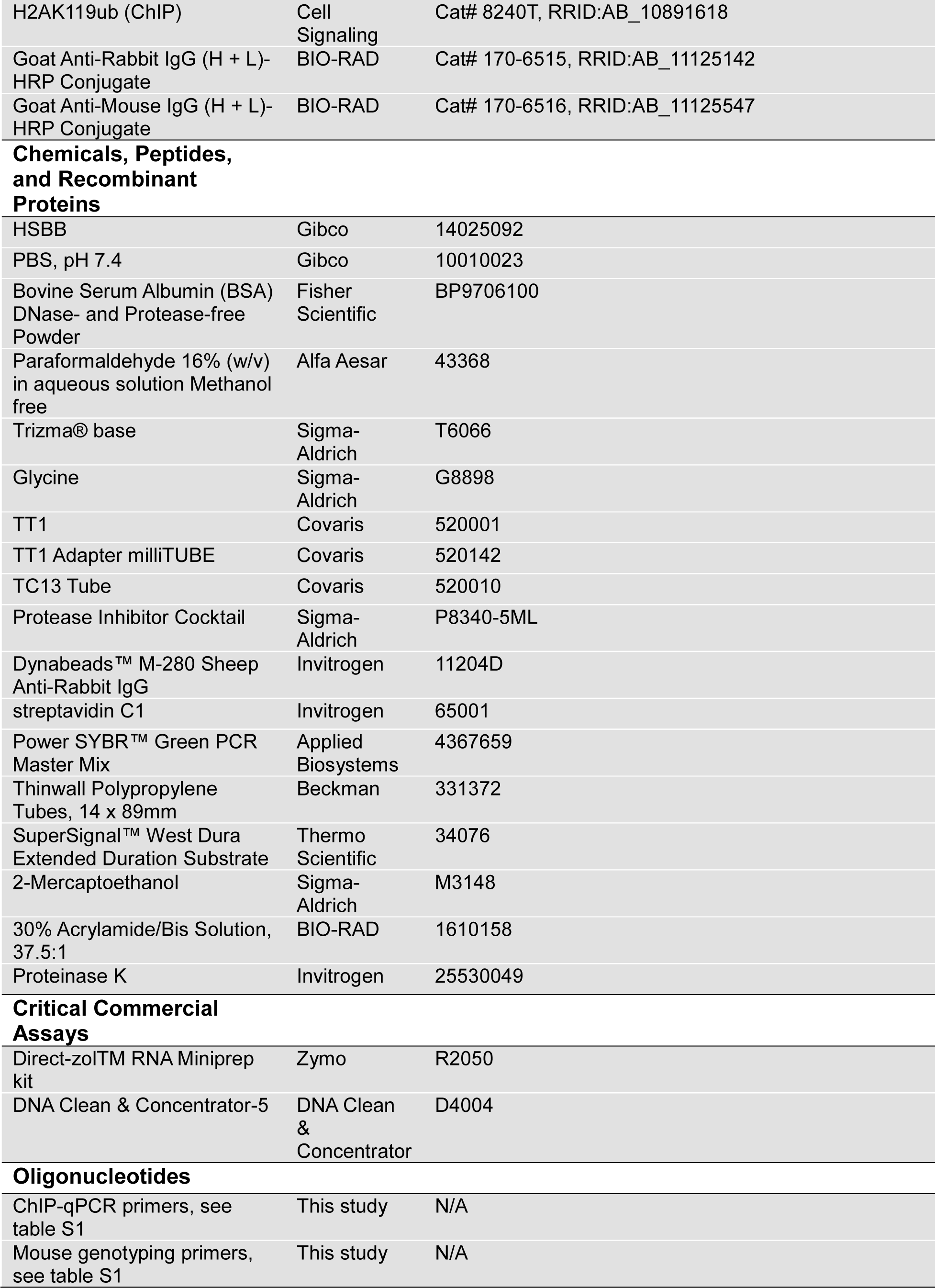
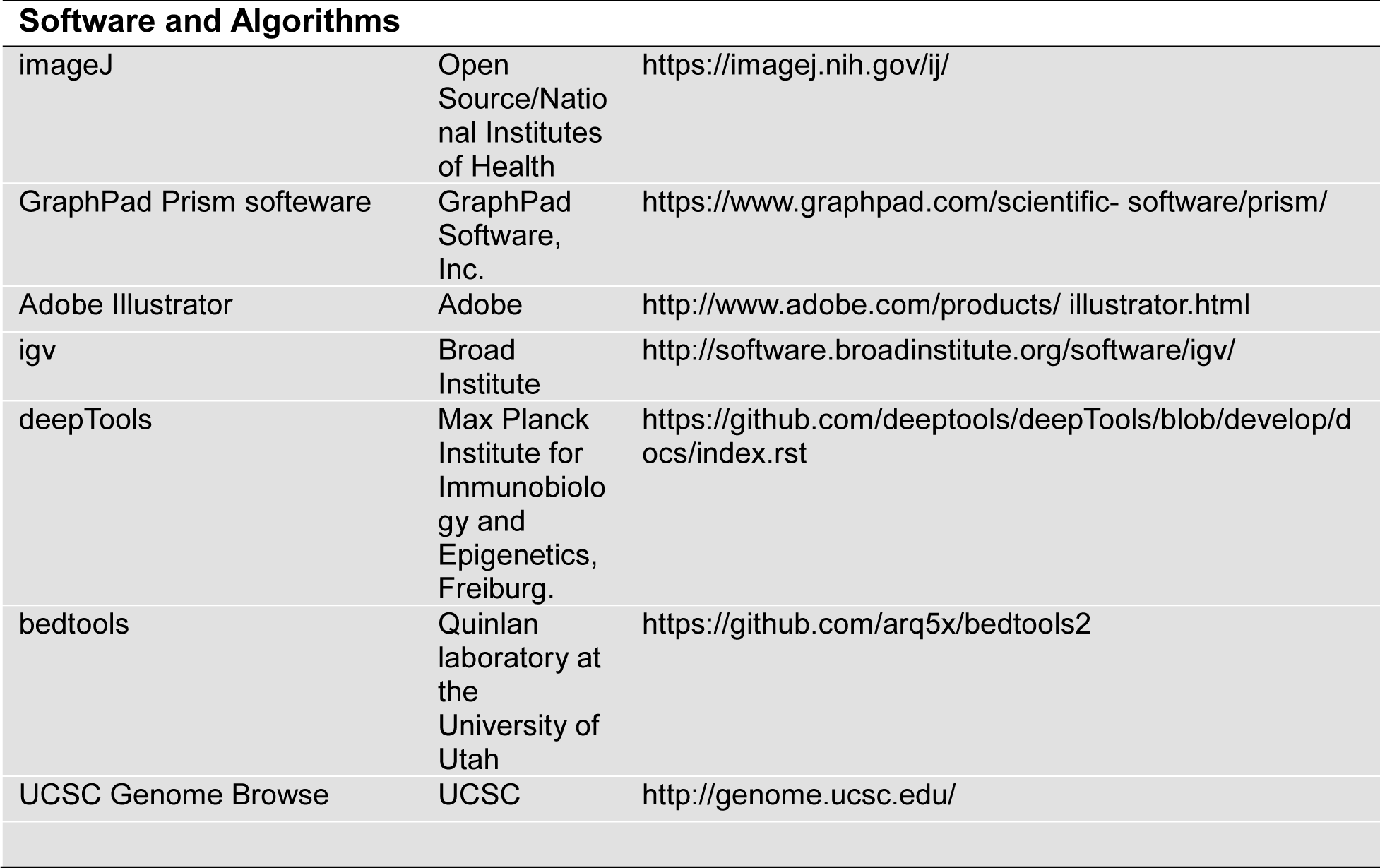

| Reagent or resource | SOURCE | IDENTIFIER |
| --- | --- | --- |
| **Antibodies** |  |  |
| Anti-SMARCA4 antibody (Western) | Abcam | Cat# ab4081, RRID:AB_304271 |
| Anti-SMARCA4 antibody (ChIP) | Abcam | Cat# ab110641, RRID:AB_10861578 |
| Anti-SMARCB1 antibody (Western) | Bethyl | Cat# A301-087A, RRID:AB_2191714 |
| Anti-SMARCC1 antibody (Western) | Proteintech | Cat# 17722-1-AP, RRID:AB_2191987 |
| Anti-SMARCC1 antibody (ChIP) | Invitrogen | Cat# PA5-30174, RRID:AB_2547648 |
| Anti-SS18 antibody (Western) | Cell Signaling | Cat# 21792, RRID:AB_2728667 |
| Anti-SS18-SSX antibody (Western and ChIP) | Cell Signaling | Cat# 72364 |
| Anti-PBRM1 antibody (Western) | Bethyl | Cat# A301-591A, RRID:AB_1078808 |
| Anti-PBRM1 antibody (ChIP) | Cell Signaling | Cat# 89123, RRID:AB_2936366 |
| Anti-ARIAD1A antibody (Western, ChIP) | Cell Signaling | Cat# 12354, RRID:AB_2637010 |
| Anti-BRD9 antibody (Western and ChIP) | Proteintech | 24785-1-AP |
| Anti-DPF2 antibody (Western and ChIP) | Abcam | Cat# ab134942, RRID:AB_2728668 |
| H3K4me1 (ChIP) | Diagenode | Cat# C15410037-50, CiteAb: pAb-037-050 |
| H3K4me3 (ChIP) | Cell Signaling | Cat# 9751S, RRID:AB_2616028 |
| H3K27ac (ChIP and HiChIP) | Abcam | Cat# ab4729, RRID:AB_2118291 |
| H3K27me3 (ChIP) | Diagenode | Cat# C15410069, CiteAb: pAb-069-050 |
| H3K36me3 (ChIP) | Abcam | Cat# ab9050, RRID:AB_306966 |
| H2AK119ub (ChIP) | Cell Signaling | Cat# 8240T, RRID:AB_10891618 |
| Goat Anti-Rabbit IgG (H + L)-HRP Conjugate | BIO-RAD | Cat# 170-6515, RRID:AB_11125142 |
| Goat Anti-Mouse IgG (H + L)-HRP Conjugate | BIO-RAD | Cat# 170-6516, RRID:AB_11125547 |
| **Chemicals, Peptides, and Recombinant Proteins** |  |  |
| HSBB | Gibco | 14025092 |
| PBS, pH 7.4 | Gibco | 10010023 |
| Bovine Serum Albumin (BSA) DNase- and Protease-free Powder | Fisher Scientific | BP9706100 |
| Paraformaldehyde 16% (w/v) in aqueous solution Methanol free | Alfa Aesar | 43368 |
| Trizma® base | Sigma-Aldrich | T6066 |
| Glycine | Sigma-Aldrich | G8898 |
| TT1 | Covaris | 520001 |
| TT1 Adapter milliTUBE | Covaris | 520142 |
| TC13 Tube | Covaris | 520010 |
| Protease Inhibitor Cocktail | Sigma-Aldrich | P8340-5ML |
| Dynabeads™ M-280 Sheep Anti-Rabbit IgG | Invitrogen | 11204D |
| streptavidin C1 | Invitrogen | 65001 |
| Power SYBR™ Green PCR Master Mix | Applied Biosystems | 4367659 |
| Thinwall Polypropylene Tubes, 14 x 89mm | Beckman | 331372 |
| SuperSignal™ West Dura Extended Duration Substrate | Thermo Scientific | 34076 |
| 2-Mercaptoethanol | Sigma-Aldrich | M3148 |
| 30% Acrylamide/Bis Solution, 37.5:1 | BIO-RAD | 1610158 |
| Proteinase K | Invitrogen | 25530049 |
| **Critical Commercial Assays** |  |  |
| Direct-zol™ RNA Miniprep kit | Zymo | R2050 |
| DNA Clean & Concentrator-5 | DNA Clean & Concentrator | D4004 |
| **Oligonucleotides** |  |  |
| ChIP-qPCR primers, see table S1 | This study | N/A |
| Mouse genotyping primers, see table S1 | This study | N/A |
| Software and Algorithms |  |  |
| imageJ | Open Source/National Institutes of Health | https://imagej.nih.gov/ij/ |
| GraphPad Prism software | GraphPad Software, Inc. | https://www.graphpad.com/scientific-software/prism/ |
| Adobe Illustrator | Adobe | http://www.adobe.com/products/illustrator.html |
| igv | Broad Institute | http://software.broadinstitute.org/software/igv/ |
| deepTools | Max Planck Institute for Immunobiology and Epigenetics, Freiburg. | https://github.com/deeptools/deepTools/blob/develop/docs/index.rst |
| bedtools | Quinlan laboratory at the University of Utah | https://github.com/arq5x/bedtools2 |
| UCSC Genome Browse | UCSC | http://genome.ucsc.edu/ |

---

**Extended Data Figure 1 (associated with Fig. 1).**
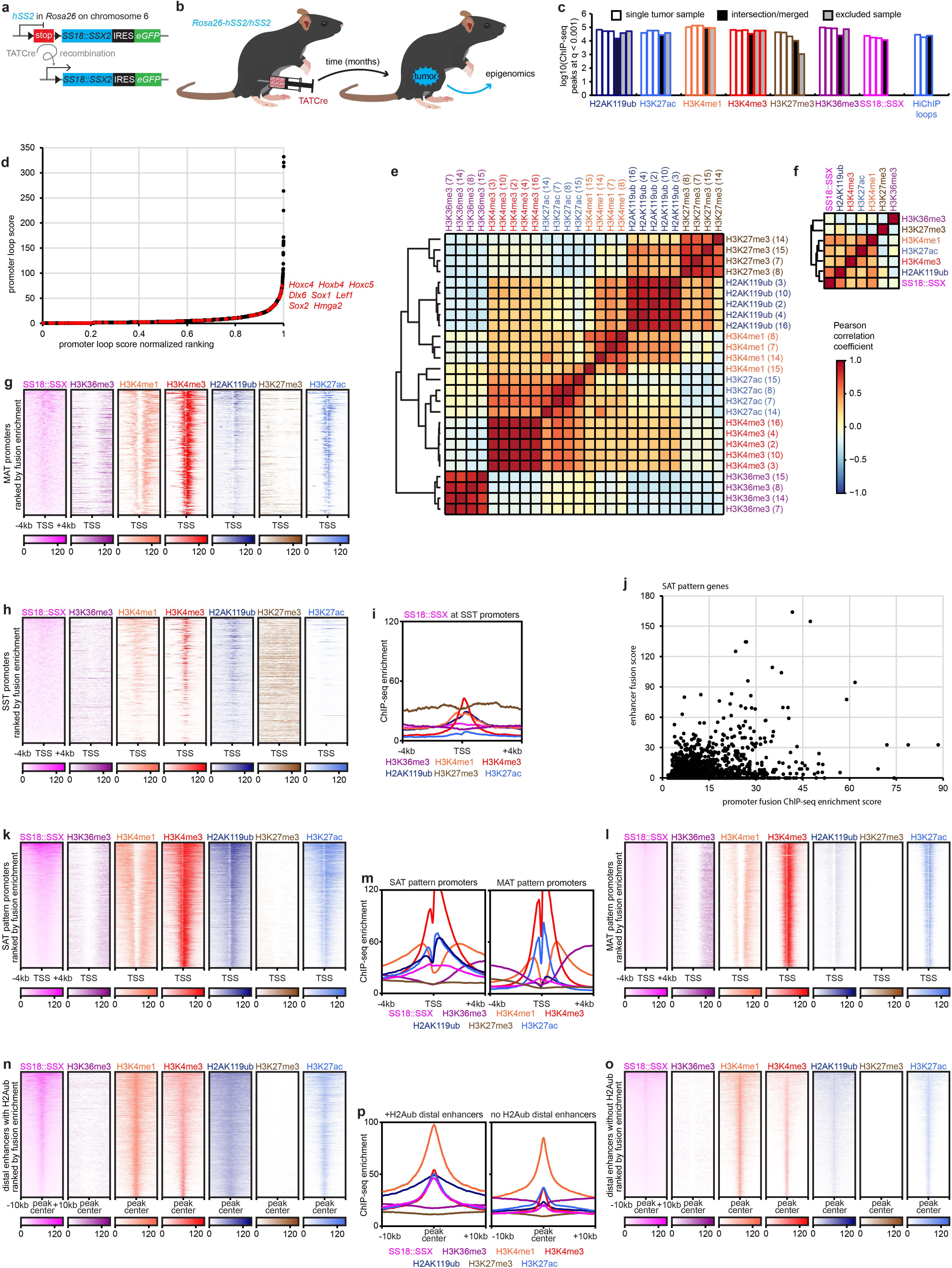
SS18::SSX distributes to the transcriptional regulatory elements of monoubiquitylated histone H2A bearing nucleosomes. (a) Schematic of the *hSS2* allele in *Rosa26*, from which the SS18::SSX2 cDNA is expressed followed Cre-mediated recombination of a floxed stop cassette. (b) Mice injected at 8 days of life in the hindlimb with TATCre develop tumors that provide the material for epigenomics assessments. (c) Peak counts after calling in MACS2 with q-value < 0.001, noting which three of each antibody’s ChIP-seq were intersected for a merged ChIP-seq peaks number shown with black filling, as well as any not included other samples. Also shown is the number of loops called for two tumors in H3K27ac HiChIP at FDR < 0.01 in Mango, then merged. (d) The H3K27ac HiChIP loops score for each looped promoter in the genome reflecting both score and a normalized ranking of the score, with SAT genes indicated in red. (e) Pearson correlation heatmap of individual samples of ChIP-seq for each indicated antibody, with the tumor numbers noted in parentheses. (f) Pearson correlation heatmap of intersection samples of each histone mark ChIP-seq with the fusion. (g) ChIP-seq enrichment heatmaps for the promoters of MAT genes. (h) ChIP-seq enrichment heatmaps and plots (i) for synovial sarcomagenesis silenced transcription genes. (j) Plot of looped distal fusion enrichment against promoter enrichment for all genes with SAT patterns in their promoters (or called peaks for H3K4me3**^+^**, H3K27ac**^+^**, H2AK119ub+, and not H3K27me3**^-^**). (k) ChIP-seq enrichment heatmaps for SAT pattern promoters and MAT pattern promoters (l) as well as enrichment plots for both (m). (n) ChIP-seq enrichment heatmaps for distal enhancers with called peaks for H3K4me1, H3K27ac, and H2AK119ub versus the same (o) for those lacking H2AK119ub called peaks and (p) enrichment plots for each.

**Extended Data Figure 2 (associated with Fig. 2).**
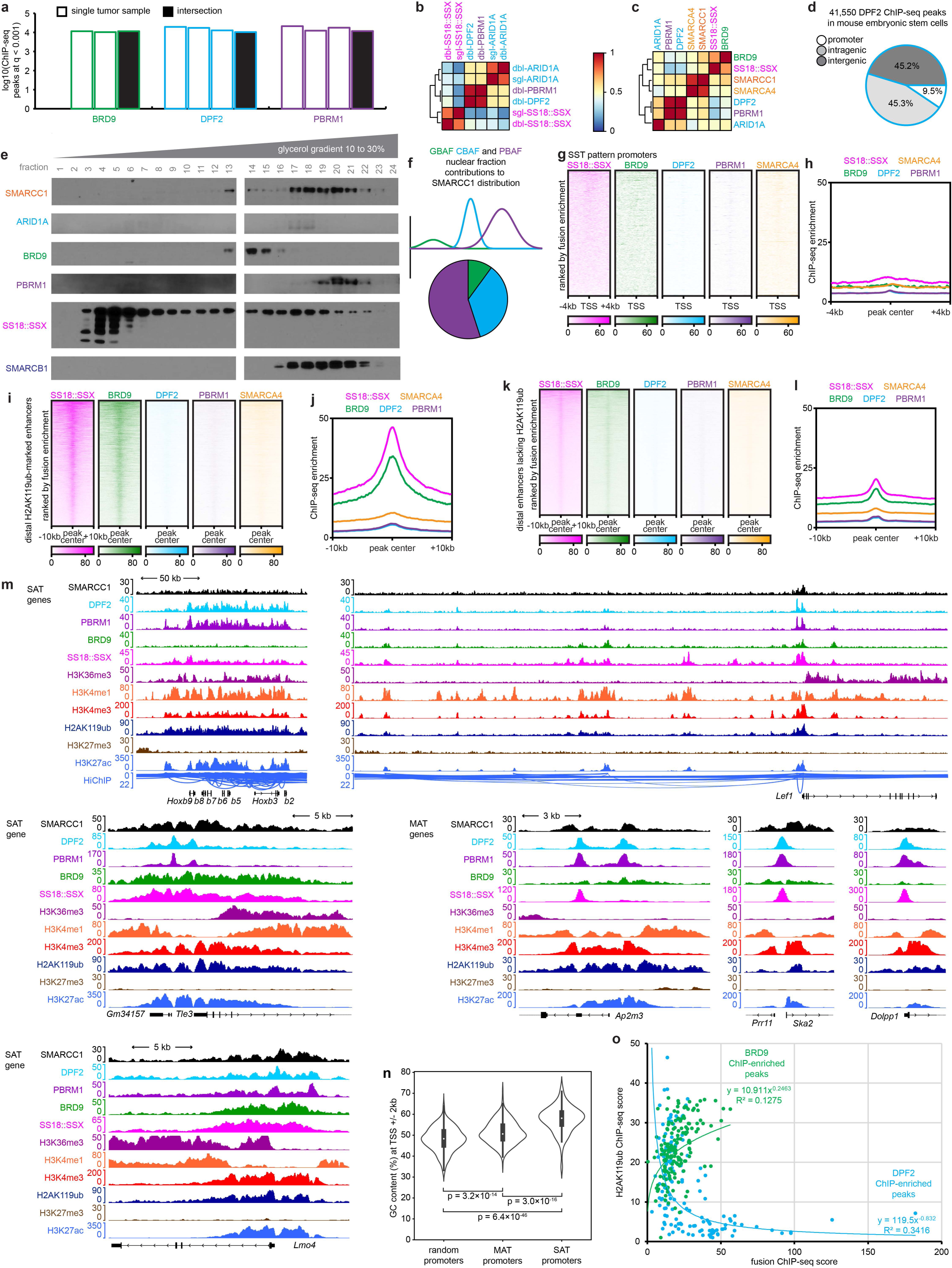
BAF family complex subtypes distribute across chromatin relative to SS18::SSX. (a) Peak counts called for BAF component ChIP-seq in individual samples, and the intersection merged peaks set. (b) Pearson correlation heatmap for individual ChIP-seq runs for the indicated antibodies after either single (sgl) or double (dbl) crosslinking. (c) Pearson correlation heatmaps of intersection merged ChIP-seq for the indicated antibodies. (d) Distribution of chromatin annotations of DPF2 ChIP-seq in mouse embryonic stem cells. (e,f) Western blots following glycerol size fractionation of nuclear extracts in a mouse SyS tumor (f) showing contributions of GBAF, CBAF, and PBAF to the overall SMARCC1 distribution calculated for each fraction. (g) ChIP-seq enrichment heatmaps for BAF components at SST gene promoters and (h) plots of the same. (i) ChIP-seq enrichment heatmaps and (j) plots for distal peaks that were determined by called peaks for H3K4me1, H3K27ac, and H2AK119ub, as well as those (k,l) lacking a called peak for H2AK119ub. (m) Example ChIP-seq enrichment tracks for SAT pattern genes with and without significant distal looped enhancers, as well as MAT genes. (n) Violin plot of the GC content in promoters of each pattern. (o) Plot of the ChIP-seq enrichment scores for fusion versus H2AK119ub, noted for those peaks that are in the top 1% of DPF2 enrichment or BRD9 enrichment, showing divergent relationships in these two populations of peaks.

**Extended Data Figure 3 (associated with Fig. 3).**
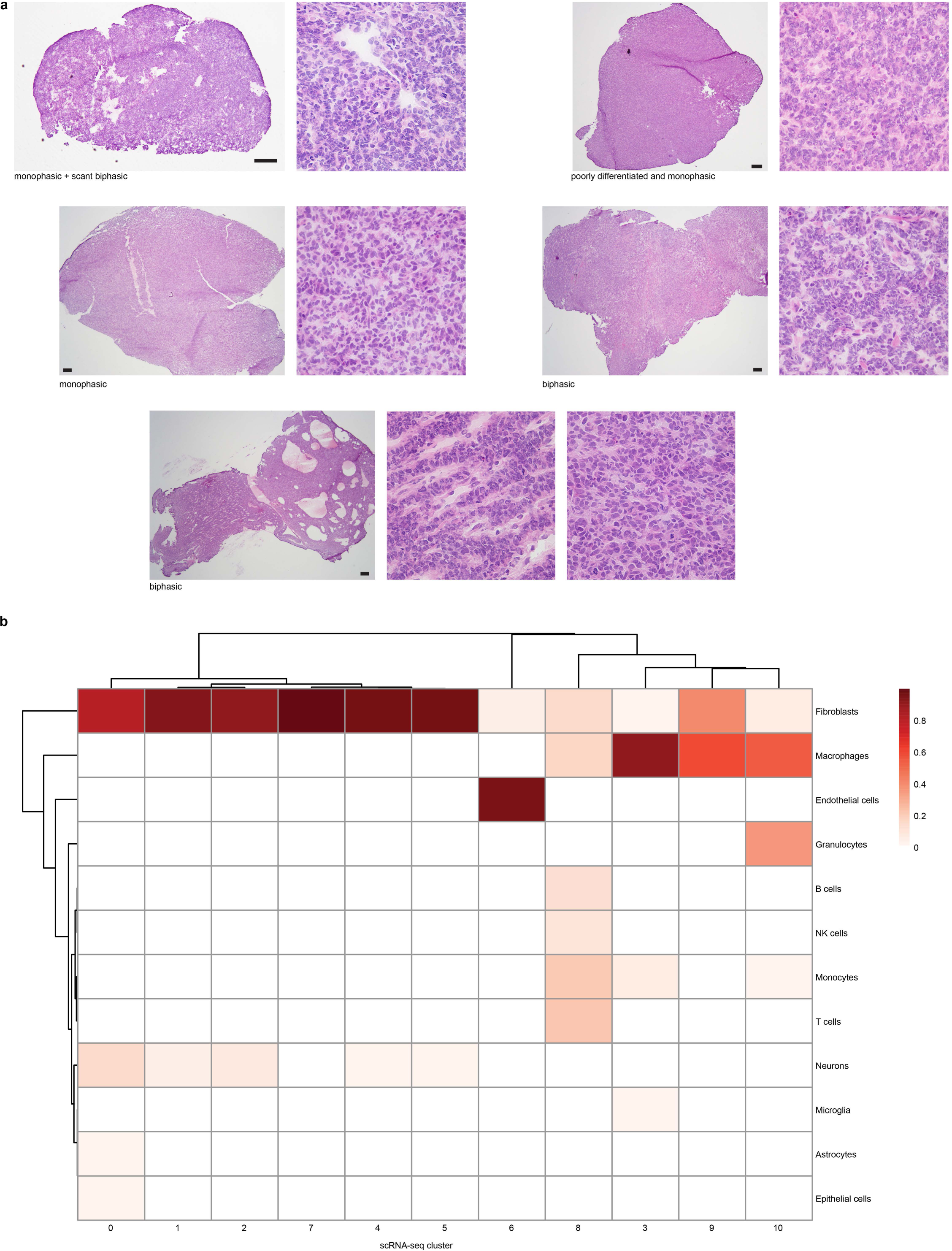
Single cell transcriptomics reveal distinct cell type clusters within synovial sarcoma tumors. (a) Representative photomicrographs showing histomorphology of snap frozen tissue sections adjacent to areas sampled by single cell RNA-seq (Magnification bars, 250µm; square panel sides, 250µm. Note that snap freezing creates artifacts resembling nuclear atypia.) (b) Identification of each of the 10 clusters by classic expression patterns in Seurat.

**Extended Data Figure 4 (associated with Fig. 3).**
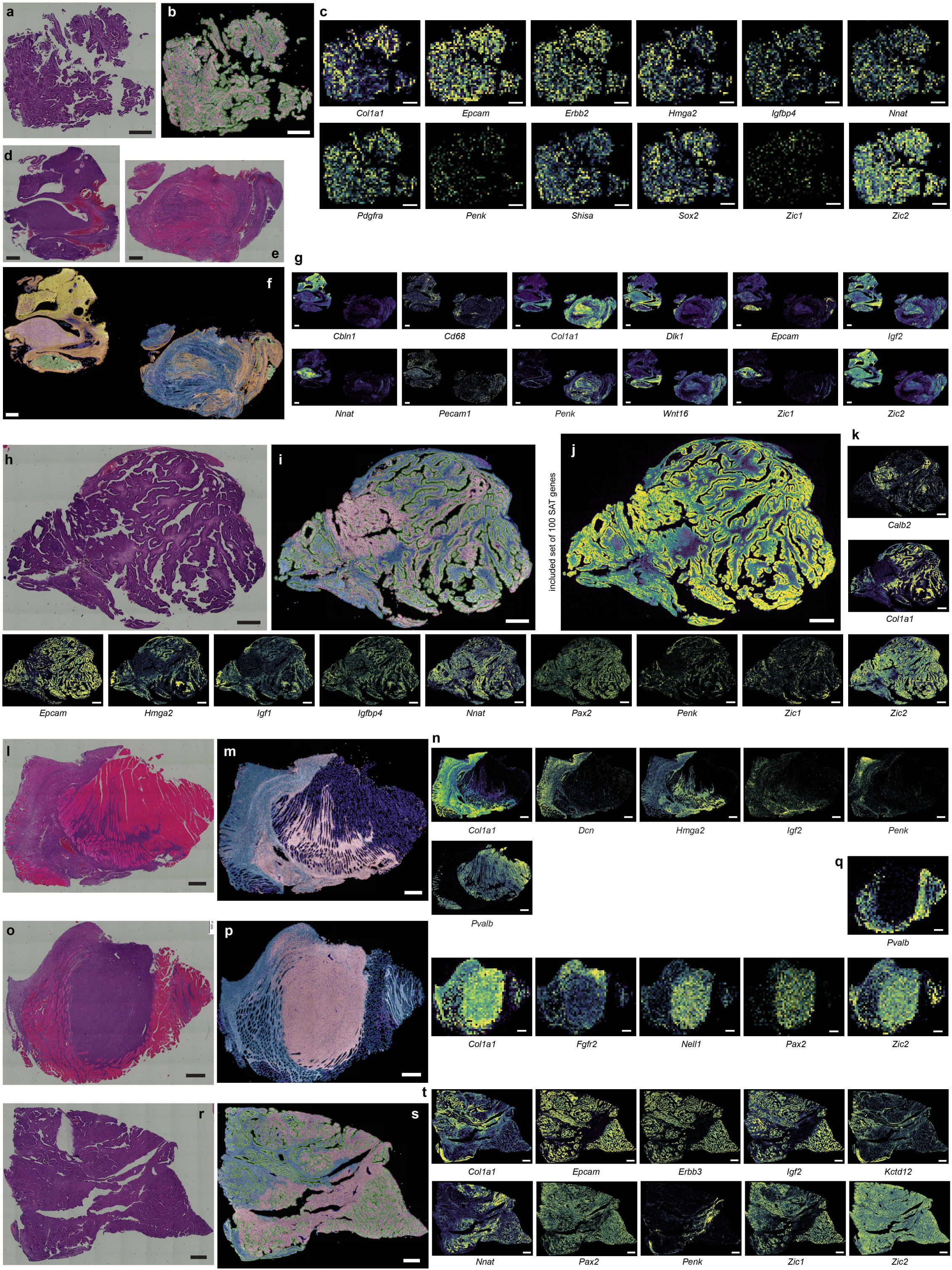
Xenium spatial transcriptomics identifies cell markers for each synovial sarcoma tissue type. (a, d, e, h, l, o, r) Photomicrographs of *hSS2* mouse tumor sections stained with hematoxylin and eosin (H&E). (b, f, I, m, p, s) Xenium In Situ pseudo-colored spatial representation of clusters of cells: monophasic (blue), epithelial and poorly differentiated (pink), glandular epithelial cells (green), endothelial cells (purple). (c, g, j, k, n, q, t) Spatial heat maps of gene expression following Xenium In Situ. (Magnification bars, 500µm).

**Extended Data Figure 5 (associated with Fig. 3).**
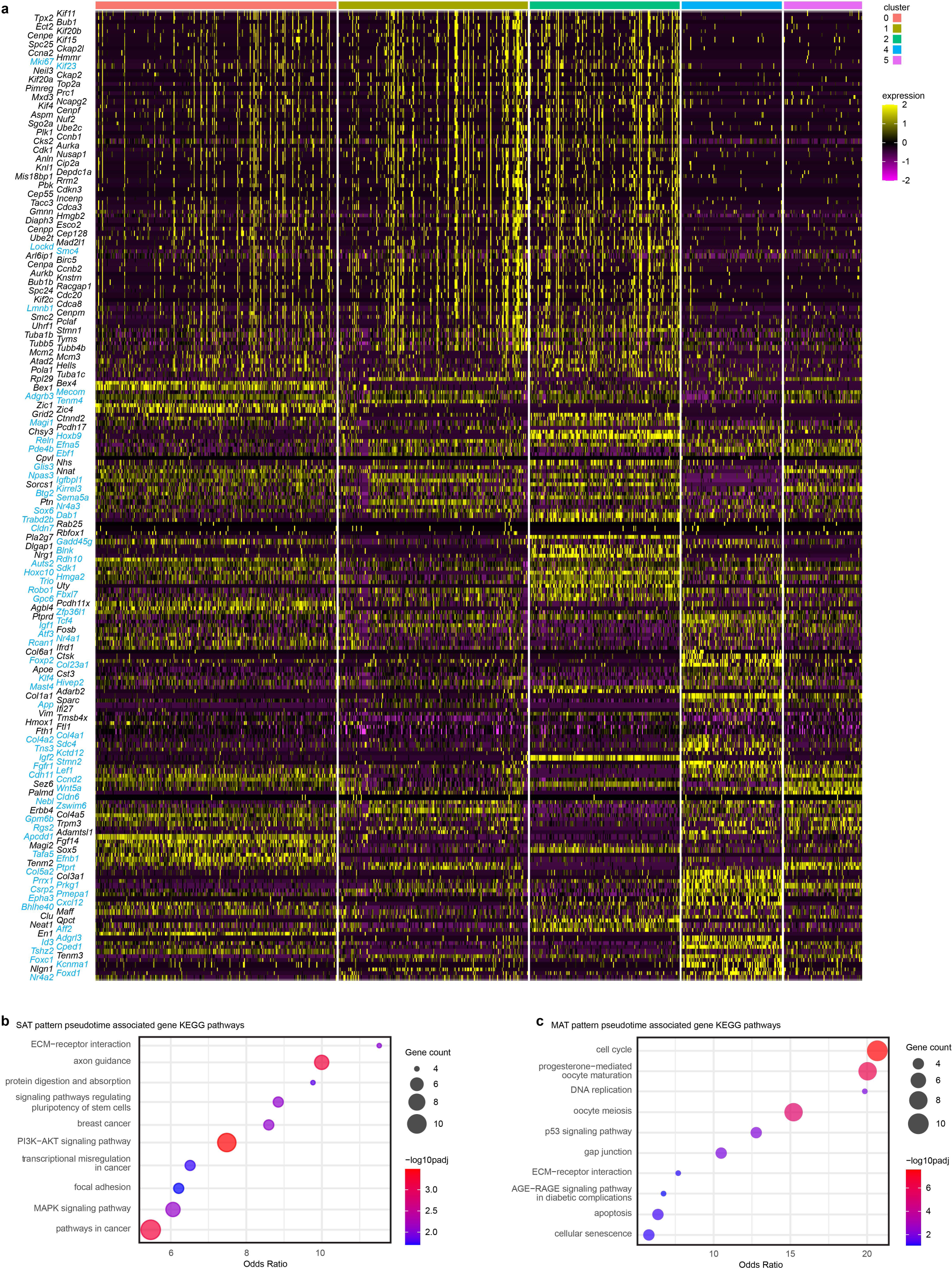
Pseudotime analysis defines gene sets associated with reprogramming. (a) Heatmap depicting gene expression profiles associated with pseudotime trajectories in tumor cell populations (SAT pattern genes, cyan text; MAT pattern genes, black text). (b) KEGG pathway analysis of SAT pattern genes and (c) MAT pattern genes, each from the pseudotime associated list.

**Extended Data Figure 6 (associated with Fig. 4).**
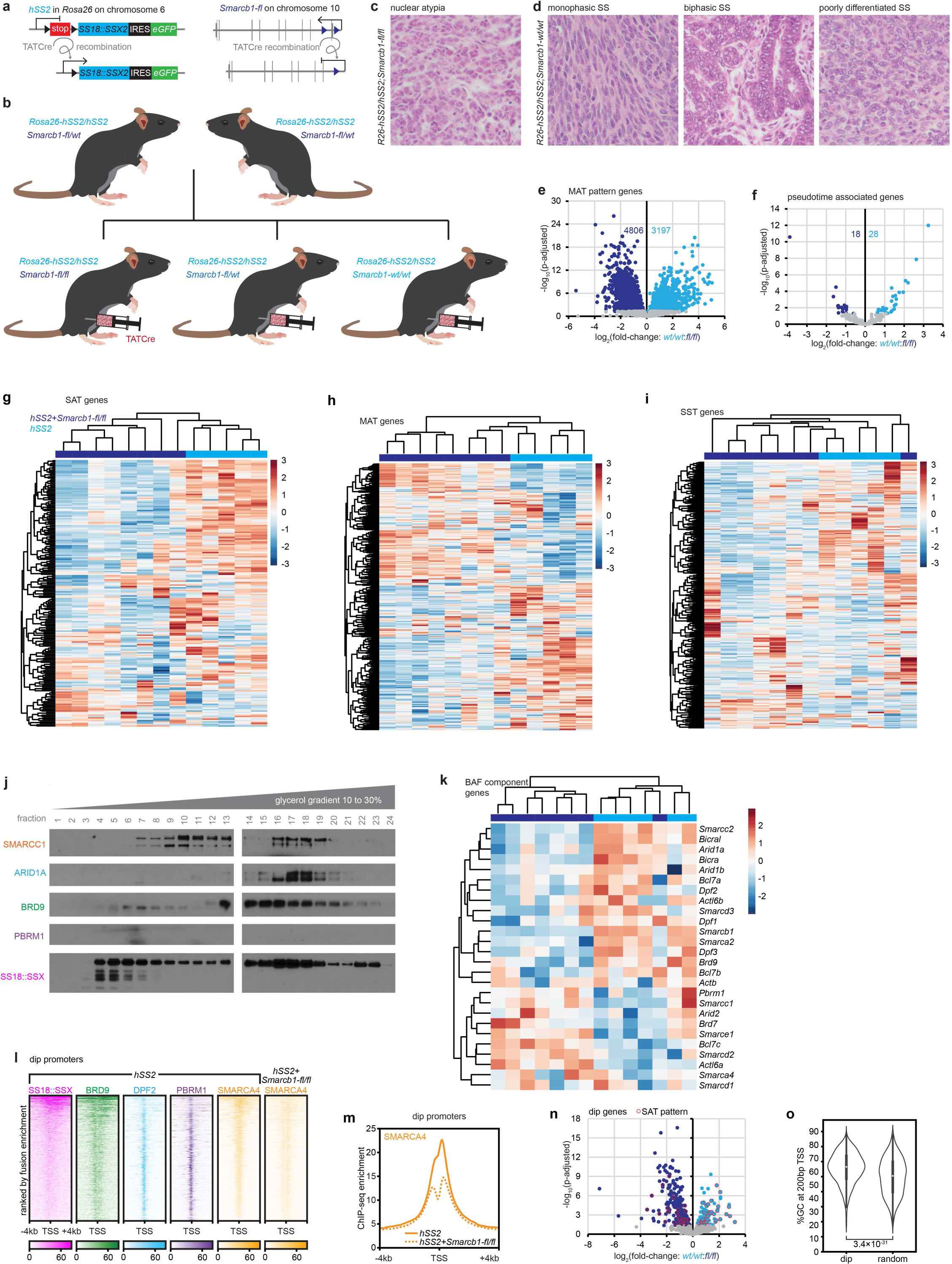
Accompanying SMARCB1 loss alters synovial sarcoma phenotypes through PBAF and CBAF impacts. (a) Schematic of the *hSS2 and Smarcb1*-floxed alleles, for simultaneous expression of SS18::SSX2 and deletion of critical *Smarcb1* exons upon Cre-mediated recombination (b) *hSS2* breeding scheme to generate littermate-controlled tumorigenesis phenotyping experiments with varied *Smarcb1* genotypes. (c) Photomicrograph example (100µm-sided square) showing nuclear atypia in an *hSS2;Smarcb1-fl/fl* tumor. (d) Photomicrograph examples (100µm-sided square) demonstrating *hSS2;Smarcb1-wt/wt* tumors with all characteristic SyS morphologies. (e) Differential expression of MAT pattern and (f) pseudotime associated genes comparing *hSS2;Smarcb1-wt/wt* and *hSS2;Smarcb1-fl/fl* tumors. (g) Expression heatmaps for SAT, (h) MAT, and (i) SST genes expressed in *hSS2;Smarcb1-wt/wt and hSS2;Smarcb1-fl/fl* tumors. (j) Western blots of BAF components following glycerol size fractionation of nuclear extracts in an *hSS2;Smarcb1-fl/fl* tumor. (k) Expression heatmaps for BAF component genes to test the depth of loss of the conditionally targeted allele and for any compensatory changes in other components. (l) Enrichment heatmaps for BAF components ChiP-seq at promoters defined as having focal loss (dip) of SMARCA4 in *hSS2*;*Smarcb1-fl/fl* tumors, as in (m). (n) Differential expression of genes defined by the dip at the promoter comparing *hSS2;Smarcb1-wt/wt* and *hSS2;Smarcb1-fl/fl* tumors. (o) Violin plots of the CG content of the 200bp surrounding the TSS with dip genes or random other promoters across the genome (p-value from Dunn test for pairwise comparisons).

**Extended Data Figure 7 (associated with Fig. 5).**
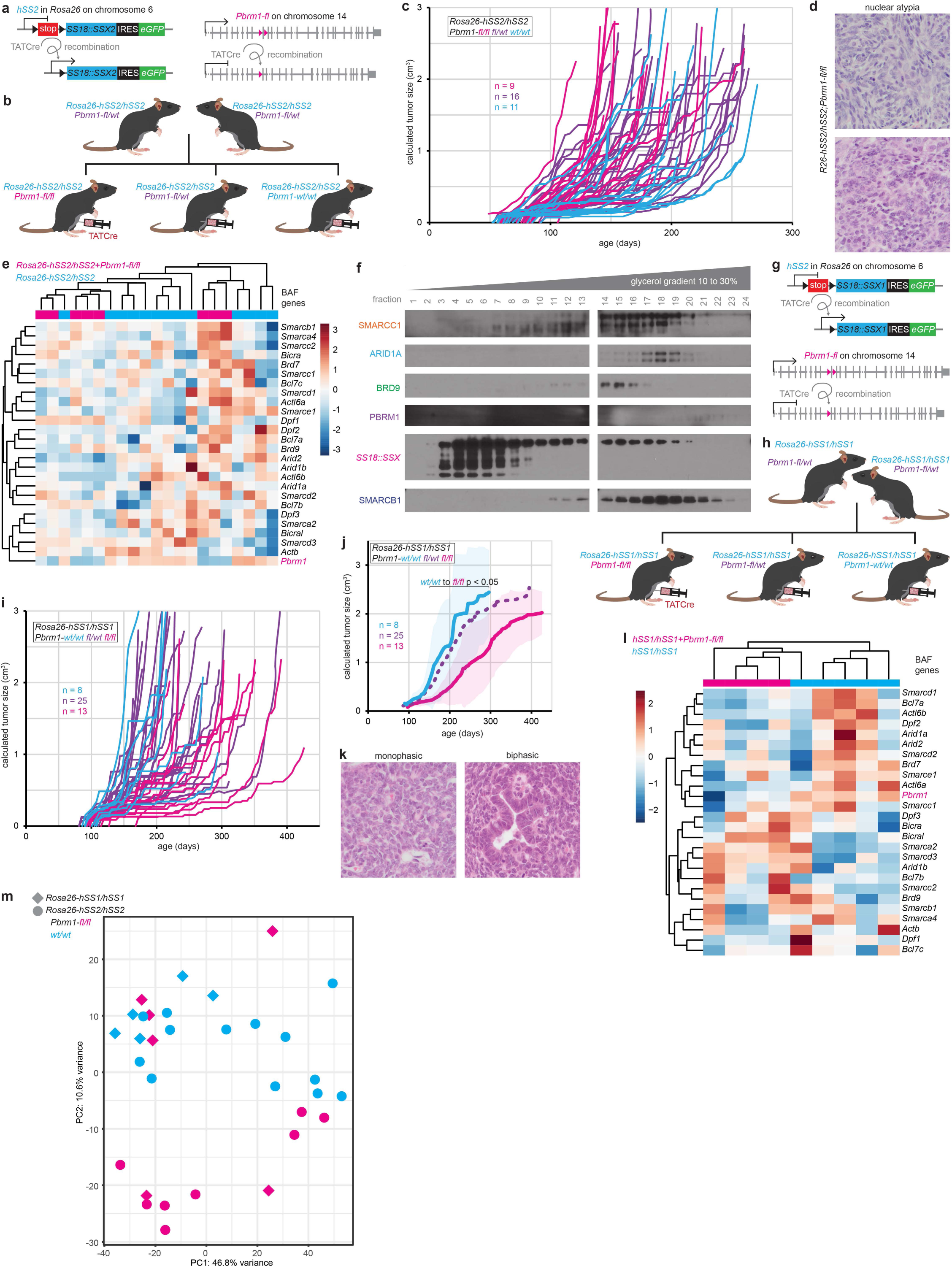
*Pbrm1* disruption blunts the SyS phenotype. (a) Schematic of the *hSS2* and *Pbrm1*-*floxed* alleles, for simultaneous expression of SS18::SSX2 and excision of *Pbrm1* exon 11 by Cre-mediated recombination. (b) *hSS2* breeding scheme to generate littermate-controlled tumorigenesis phenotyping experiments with varied *Pbrm1* genotypes. (c) Individual tumor growth trajectories for cohorts of hSS2 mice with varied *Pbrm1* genotypes. (d) Photomicrograph examples demonstrating modest nuclear atypia in *hSS2;Pbrm1-fl/fl* tumors. (e) Expression heatmap of BAF component genes in *hSS2* tumors with or without Pbrm1 disruption. (f) Western blots for BAF components after glycerol gradients of nuclear fractions of a *hSS2;Pbrm1-fl/fl* tumor. (g) Schematic of *hSS1* allele and the *Pbrm1*-*floxed* allele and the breeding scheme (h) to generate littermate-controlled tumor phenotyping experiments. (i) Individual growth curves and aggregate (j) curves demonstrating mean tumor size (shading indicates ± standard deviations, p-values from two-tailed, heteroscedastic *t* tests) in *hSS1* mice injected with TATCre at day 8 of life comparing littermates with varied *Pbrm1-fl* genotypes. (k) Photomicrograph examples demonstrating retained SyS phenotypes without nuclear atypia in the *hSS1;Pbrm1-fl/fl* tumors which develop more slowly than tumors in *hSS2;Pbrm1-fl/fl* mice. (l) Expression heatmap for bulk RNA-seq of BAF components among *hSS1* tumors arising in mice with homozygous floxed or wildtype *Pbrm1*, demonstrating only subtle reduction in *Pbrm1* in the homozygous floxed tumors, suggesting heterozygous recombination of the floxed alleles in the cells that gave rise to these tumors. (m) Principal Component Analysis plot in two dimensions from bulk RNA-seq of whole tumor transcriptomes in *hSS1;Pbrm1-fl/fl*, and *hSS2;Pbrm1-fl/fl*, mice.

**Extended Data Figure 8 (associated with Fig. 6).**
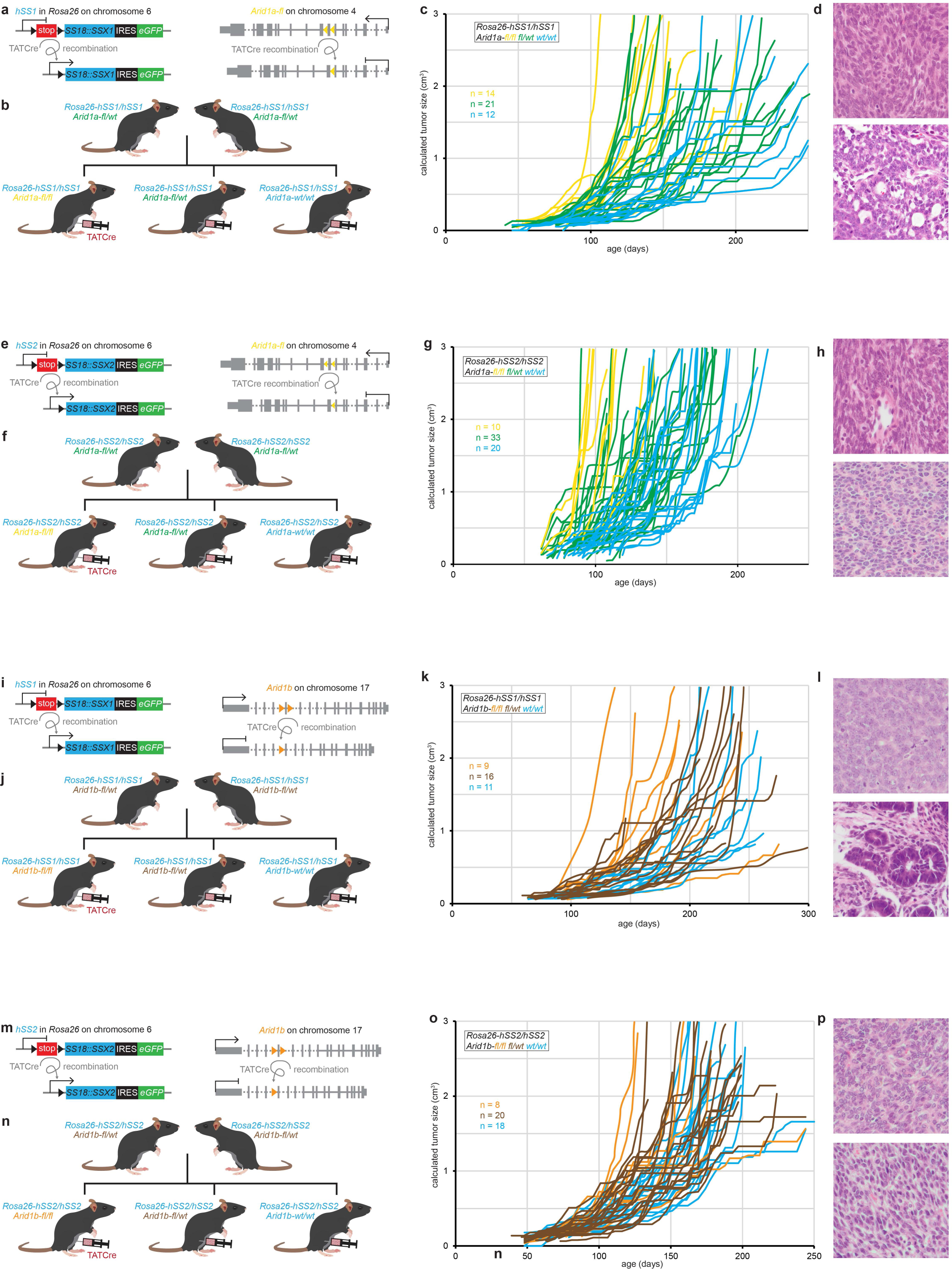
CBAF component disruptions enhance synovial sarcomagenesis and retain the SyS phenotype. (a) Schematic of *hSS1* and *Arid1a-floxed* alleles, for simultaneous expression of SS18::SSX1 and excision of *Arid1a* exon 8 by Cre-mediated recombination. (b) Breeding scheme for generating littermate-controlled cohorts of *hSS1* mice bearing variable Arid1a genotypes. (c) Individual growth curves for *hSS1* tumors with variable *Arid1a* genotypes, as well as (d) representative photomicrographs, showing retained synovial sarcoma histomorphology. (e-h) Alleles, breeding scheme, tumor growth curves, and representative photomicrographs for *hSS2* phenotyping experiments with varied Arid1a genotypes. (i-l) *hSS1* and (m-o) *hSS2* mice with *Arid1b* genotypes. The *Arid1b-floxed* allele deletes exon 5 via Cre-mediated recombination.

**Extended Data Figure 9 (associated with Fig. 6).**
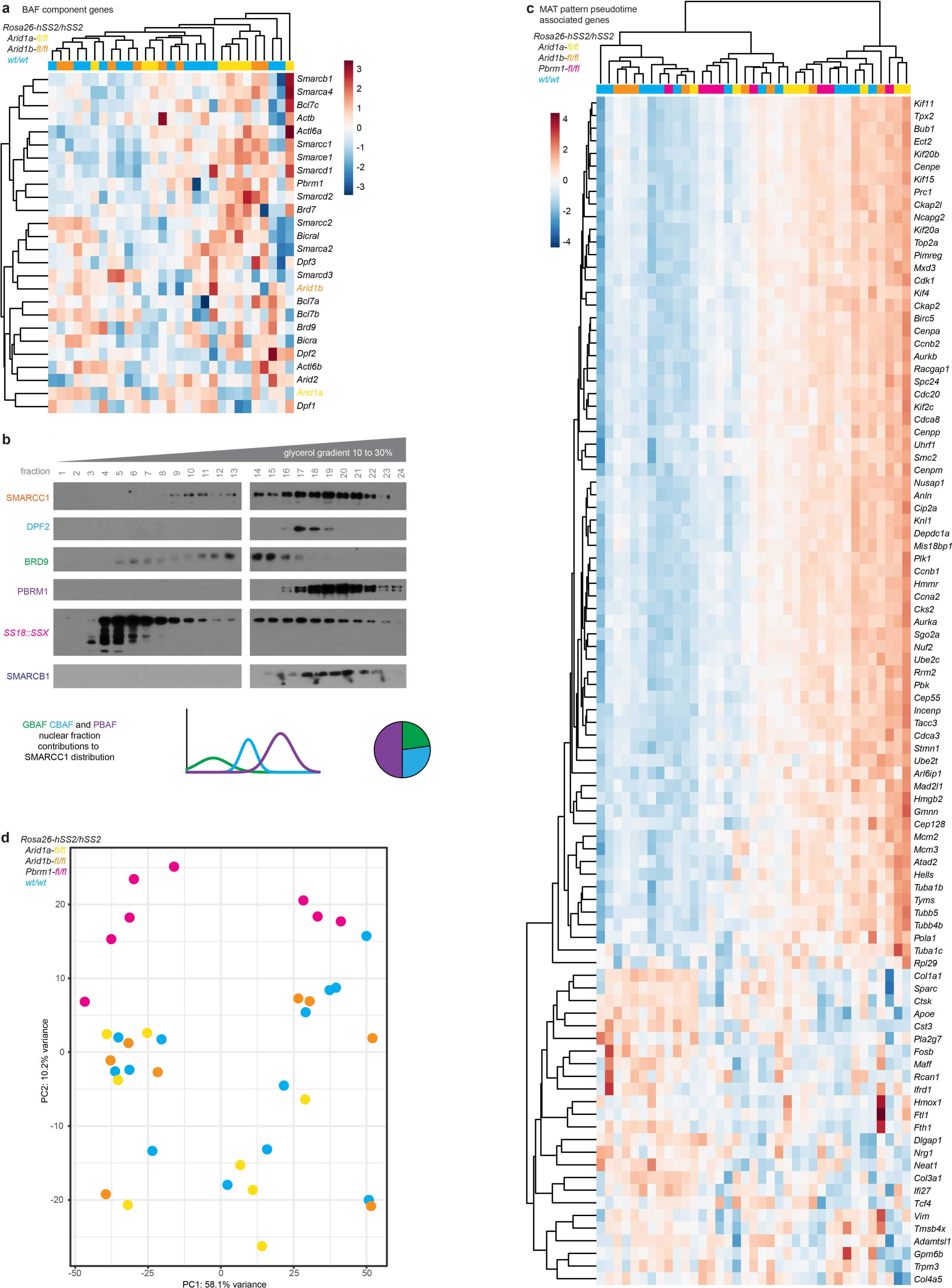
CBAF component disruptions are not compensated, driving further loss of SST genes. (a) Expression heatmap of BAF component genes showing a lack of paralog gene upregulation when either *Arid1a* or *Arid1b* is disrupted in the presence of the other. (b) Western blots of BAF components following glycerol size fractionation of nuclear extracts, showing contributions of CBAF, GBAP and PBAF to the overall SMARCC1 distribution calculated for each fraction. (c) Expression heatmap for non-SAT pattern genes identified as associated with pseudotime trajectories, tested by bulk RNA-seq of *hSS2* tumors with varied *Arid1a*, *Arid1b*, and *Pbrm1* genotypes. (d) PCA plot of whole transcriptomes for *hSS2* tumors with varied *Arid1a*, *Arid1b*, and *Pbrm1* genotypes.

**Extended Data Figure 10 (associated with Fig. 6).**
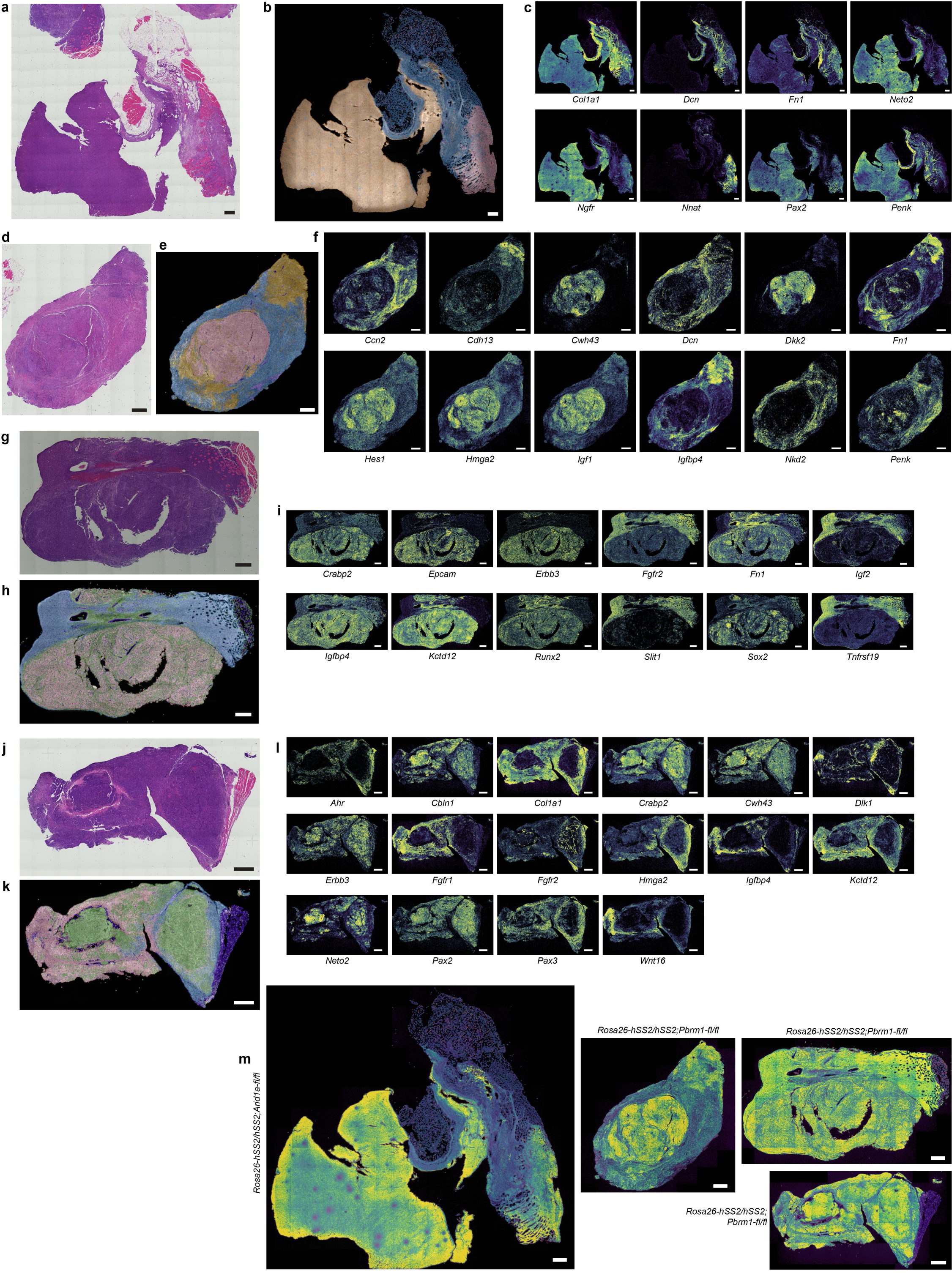
CBAF disruption generates mostly poorly differentiated tumors with strong regprogramming gene signatures, while PBRM1 disruption alters the expression of many other genes, changing the relationships between specific SAT genes and certain cellular phenotypes. (a) hematoxylin and eosin (H&E) photomicrograph of a *Rosa26-hSS2/hSS2;Arid1a-fl/fl* tumor with (b) Xenium clustering annotation and (c) expression heatmaps for individual genes. (d, g, j) H&E photomicrographs of *Rosa26-hSS2/hSS2;Pbrm1-fl/fl* tumors with Xenium clustering (e, h, k) and expression heatmaps for individual genes (f, I, l). (m) Heatmaps for expression of a 100 SAT gene set included on the Xenium. (All magnification bars are 500µm in length).

